# *Thalassoporum longitrichum* sp. nov., a marine epizoic cyanobacterium with anti-inflammatory potential, and the taxonomic reassessment of *Limnothrix* Meffert

**DOI:** 10.64898/2026.03.11.711011

**Authors:** C. Tenorio Rodas, G. S. Hentschke, F. Oliveira, G. Lopes, A. Duarte, J. Morone, A. Campos, V. Vasconcelos

## Abstract

The strain LEGE 10371, isolated from the surface of a marine sponge at Praia da Memória, Portugal, was characterized as a new *Thalassoporum* species (Pseudanabaenales) using a polyphasic approach that included 16S rRNA gene phylogenetic analysis (Maximum Likelihood and Bayesian Inference), 16S-23S ITS secondary structures, p-distance calculations, MALDI-TOF MS profiling, and morphological analysis by optical and scanning electron microscopy, as well as ecological and biochemical characterization. Phylogenetically, LEGE 10371 clustered within the *Thalassoporum* clade, however distant from the other existent species of the genus. The p-distance analysis revealed low sequence identity with other *Thalassoporum* species, with a maximum value of 97.2% to *Th. komareki*. The MALDI-TOF profile displayed high-intensity peaks at approximately 3,000, 4,000, 6,000 and 8,000 m/z, representing strong candidates for diagnostic markers of the new species. Morphologically, the new species differ from the other species of the genus by presenting trichomes with more than 10 cells and lack of aerotopes. Biocompatibility of the fractions was evaluated in HaCaT keratinocytes, showing no cytotoxic effects at most tested concentrations. PCR screening targeting *mcyE*, *sxtG*, *anaC*, and *cyrA* confirmed the absence of the genetic potential for the production of major cyanotoxins. Chemical characterization revealed a pigment-rich profile dominated by chlorophyll-a and carotenoids, including β-carotene, zeaxanthin, lutein, and mixoxanthophyll. Bioactivity assays showed superoxide anion radical scavenging by the aqueous fraction (IC₂₅ ≈ 0.042–0.045 mg mL⁻¹), strong nitric oxide radical scavenging by the acetonic fraction (IC₅₀ = 0.045 mg mL⁻¹), and lipoxygenase inhibition (∼41%, for a fraction concentration of 0.25 mg mL⁻), suggesting a potential contribution of these fractions to modulate inflammation-related pathways. Additionally to this results, the polyphasic analysis permitted to confirm previous data that *Pseudanabaena* and *Limnothrix* represent the same generic entity. Both genera clustered together, presented high 16S rRNA gene identity (up to 99.9%) and share the same morphological and ecological features. Consequently, we formally proposed the synonimization of *Limnothrix* into *Pseudanabaena*.

## Introduction

Cyanobacteria are photosynthetic prokaryotes with an estimated origin of approximately 3.5 billion years. They play a fundamental ecological role in aquatic and global ecosystems, acting as primary producers, nitrogen fixers, and modulators of biogeochemical cycles (Whitton, 2012).

The classification of cyanobacteria has long posed a significant challenge due to their extensive diversity and the continuous advancement of analytical methods. Early systematic efforts, made by Bornet & Flahault (1886, 1888), Gomont, (1892), and later by Komárek & Anagnostidis (1986, 1988,2005) allowed the description and revision of numerous species based primarily on botanical classical morphological characters. Later, (Komárek et al., 2014) conducted a comprehensive taxonomic revision using a polyphasic approach that integrated morphology, genetics, ecology, and biochemistry, thereby redefining the internal hierarchy of cyanobacteria. More recently, Strunecký et al. (2023) integrated phylogenomic analyses to cyanobacterial systematics and defined twenty orders in the class Cyanophyceae. Within this updated framework, the family Pseudanabaenaceae has undergone successive reassignments, from Oscillatoriales (Komárek & Anagnostidis, 2005) to Synechococcales (Komárek et al., 2014), and most recently to Pseudanabaenales (Strunecký et al., 2023).

According to (Komárek et al., 2014), Pseudanabaenaceae originally comprised only two genera: *Pseudanabaena* Lauterborn and *Limnothrix* Meffert. In 2021, the type species *Thalassoporum komarekii* Konstantinou & Gkelis was provisionally assigned to this family based on its morphological similarity to thin filamentous pseudanabaenacean taxa (Konstantinou et al., 2021). Phylogenetic analyses subsequently demonstrated that *Thalassoporum* forms a well-supported lineage distinct from *Pseudanabaena* sensu stricto and related genera (Konstantinou et al., 2021), and the family Thalassoporaceae was proposed to reflect this evolutionary separation (Strunecký et al., 2023). More recently, a second species, *Th. mexicanum* Saraf et al., was described for the genus (Saraf et al., 2025)

For the Pseudanabaenales, in addition to these aforementioned genera, *Tumidithrix elongata* Luz et al. was described from freshwater habitats in the Azores (Portugal) (Luz et al., 2023). Within the order, *Pseudanabaena, Limnothrix, Tumidithrix* and *Thalassoporum,* are characterized by unbranched filaments composed of cylindrical cells typically <4 μm wide, lacking akinetes and true differentiated thalli. Their cells lack thick envelopes and contain parietal thylakoids, traits historically considered diagnostic for *Pseudanabaenaceae.*(Anagnostidis & Komárek, 1988; Komárek & Anagnostidis, 2005; Komárek et al., 2014; Konstantinou et al., 2021; Luz et al., 2023). However, two coccoid genera, *Portococcus* Strunecký et al. and *Pseudanabaenococcus* Strunecký et al., were recently described for the Pseudanabaenales, expanding the circumscription of the order to include not only thin filamentous forms (Strunecký et al., 2023).

Among these genera, the similarities among *Pseudanabaena and Limnothrix* are remarkable. Ecologically, they exhibit highly opportunistic strategies, tolerating broad gradients of light and nutrient availability. They occur in fresh, brackish, and coastal waters, where they frequently form dominant populations under eutrophic conditions, often contributing to recurrent blooms (Napiórkowska-Krzebietke et al., 2023; Paerl & Otten, 2013; Rojo & Alvarez Cobelas, 1994).

Morphologically, *Pseudanabaena* and *Limnothrix* have traditionally been distinguished based on trichome organization and the occurrence of aerotopes (Komárek & Anagnostidis, 2005). In this framework, *Pseudanabaena* is characterized by solitary trichomes or fine mats, usually constricted at the cross-walls and exhibiting facultative aerotopes, whereas *Limnothrix* is defined by solitary trichomes generally lacking constrictions, with aerotopes typically present during the vegetative stage. However, Komárek & Anagnostidis, (2005) already emphasized that clear boundaries between these genera are lacking under the morphological concept, as numerous *Pseudanabaena* species had been reported to display features previously considered diagnostic of *Limnothrix*. For example, *P. galeata* SAG 13.83 and *P. pruinosa*, had been reported to possess aerotopes (Aleksovski et al., 2024), a typical *Limnothrix* character, reinforcing the need for a revision of these genera to achieve a more stable and phylogenetically meaningful taxonomic framework.

Besides these morphological similarities, phylogenetically, *Pseudanabaena* and *Limnothrix* are strongly related. The reference strain for the type species of *Limnothrix*, *L. redekei* PCC 9416, has a very short 16S rRNA gene sequence, which shows 100% similarity with other strains such as CCAP 1443/1 and NIVA-CYA 227/1, both assigned to the *Pseudanabaena* clade (Akagha et al., 2019; Aleksovski et al., 2024; Luz et al., 2023).

In agreement with the morphological and phylogenetic similarities, studies have revealed that the 16S rRNA gene divergence between *Limnothrix* and *Pseudanabaena* is remarkably low. The reference strain *L. redekei* PCC 9416 shares more than 98% of 16S rRNA gene sequence identity with *Pseudanabaena* strains, challenging the traditional morphological boundaries between the two genera (Acinas et al., 2008; Akagha et al., 2019). Identity values above 94.5% are strong indicators of affiliation with the same genus (Yarza et al., 2014)

Taken together, these evidences indicate that *Pseudanabaena* and *Limnothrix* are indistinguishable (Akagha et al., 2019). Other morphologically related strains designated as *L. limentica, L. rosea*, and *L. redekei* cluster at divergent positions in Nodosilineales (Persinemataceae) and Chroococcales (Geminocystaceae), and should be revised (Akagha et al., 2019; Gkelis et al., 2005; Strunecký et al., 2023; Zhu et al., 2012).

Beyond the taxonomic complexity of this lineage, members of the Pseudanabaenales have increasingly attracted attention due to their emerging biotechnological potential. Cyanobacteria are characterized by a complex and diverse pigment composition, including chlorophyll a, carotenoids and phycobiliproteins, which play fundamental roles in photosynthesis and photoprotection but have also been associated with relevant bioactive properties (Komárek et al., 2014). In particular, carotenoids and phycobiliproteins have been linked to antioxidant capacity through free radical scavenging mechanisms and redox modulation, as well as to the regulation of inflammatory pathways (Duarte et al., 2026; Rodrigues et al., 2024). The functional relevance of these pigments extends beyond simple reactive species neutralization, as oxidative stress and inflammation are closely interconnected biological processes.

Recent studies have demonstrated that pigment-rich cyanobacterial extracts are capable of scavenging relevant physiologic free radicals while simultaneously modulating key pro-inflammatory enzymes such as inducible nitric oxide synthase (iNOS) and lipoxygenase (LOX), leading to the reduction of nitric oxide production and attenuation of inflammatory signaling cascades (Rodrigues et al., 2024). These findings support the concept that cyanobacterial pigments may exert dual antioxidant and anti-inflammatory effects by interfering with oxidative stress–mediated pathways. Moreover, the polyphasic taxonomic framework currently applied to cyanobacteria has highlighted the metabolic diversity present within filamentous lineages, further supporting their potential as sources of bioactive secondary metabolites (Komárek et al., 2014;Leão et al., 2012)

Although specific bioactivity studies focusing exclusively on *Pseudanabaena*, *Limnothrix* and *Thalassoporum* remain limited, their close phylogenetic relationship to other pigment-producing filamentous cyanobacteria suggests comparable biochemical potential. Marine taxa have demonstrated distinctive pigment profiles linked to functional activities including antioxidant and anti-inflammatory effects (Duarte et al., 2026; Favas et al., 2022; Morone et al., 2022). The demonstrated capacity of cyanobacterial pigments to interfere with oxidative stress–driven inflammatory cascades reinforces the hypothesis that members of this lineage may represent promising yet underexplored sources of anti-inflammatory compounds.

In this context, this study aims to clarify the long-standing ambiguity between *Pseudanabaena* and *Limnothrix*, and adds new diversity to the group with the description of a new species of *Thalassoporum*. To further characterize this species, its chemical profile was established by high performance liquid chromatography (HPLC), and its bioactive metabolites were explored for their potential to interact with important inflammatory mediators.

## Material and Methods

### Sampling

Strain LEGE 10371 was isolated in 2010 from the surface of a marine sponge collected in the intertidal zone of Praia da Memória, Portugal (41.23119° N, 8.72175° W). The strain was isolated by micromanipulation using Z8 supplemented with 25 g L^−1^ (Z8_25_) of synthetic sea salt (Tropic Marine, Berlin, Germany) and 10 µg mL^−1^ of vitamin B12 (Kotai, 1972) and was deposited at the Blue Biotechnology and Ecotoxicology Culture Collection (LEGE-CC) at CIIMAR. The strain is currently maintained in the same medium under the following conditions: 19°C under a 12:12 h light:dark cycle (25 μmol photons m^−2^ s^−1^). An aliquot of the strain was lyophilized and is preserved in the University of Porto Herbarium under the code PO-T5292.

### Morphological analysis and Transmission electron microscopy (TEM)

LEGE 10371 was observed, microphotographed, and analyzed using MOTIC Panthera light microscope, with camera and dedicated software (MOTIC Europe, Barcelona, Spain) and Leica DMLB light microscope, also equipped with camera. The cells measurements were carried out at various positions of the slide’s preparations, totalizing 20 to 30 measurements per strain. To visualize the mucilaginous envelopes, China Ink was used for contrast.

SEM analysis was used to examine cell surface morphology. Cells were harvested by centrifugation and fixed overnight (12 h) at 4 ◦C in a final solution of 2% glutaraldehyde (for electron microscopy, VWR BDH Chemicals, Prague, Czech Republic) prepared in 50mM sodium cacodylate buffer, pH 7.4 (Electron Microscopy Sciences, PA, USA). After being postfixed overnight, cells were washed once in a double-strength sodium cacodylate buffer. Dehydration was performed through a graded ethanol series in deionized water (v/v) of 25%, 50%, 75%, and 100%, with each step lasting 5 min (Ethanol Absolute, Molecular Biology Grade, Fisher BioReagents, Loughborough, United Kingdom). Between steps, cells were collected by centrifugation at 11,000× g for 1 min at room temperature (Micro Star 17R, VWR, Radnor, PA, USA). In the final step, 500 μL of 100% ethanol was added to fully submerge the sample. Critical-point drying was used (CPD 7501, Polaron Range, Warsaw, Poland) to preserve the sample’s ultrastructure. The sample was coated with a gold/palladium (Au/Pd) thin film in adhesive carbon tape by sputtering, using the SPI Module Sputter Coater equipment for 80 s and a 15 mA current. SEM/EDS analysis was performed using a high-resolution (Schottky) Environmental Scanning Electron Microscope with X-Ray Microanalysis and Electron Backscattered Diffraction analysis: FEI Quanta 400 FEG ESEM/EDAX Genesis X4M. Images were taken with the SEM software version 4.1.10.2127 at an acceleration voltage of 15kV with the Everhart–Thornley detector (ETD) for secondary electrons (SE) and Backscattered Electron Detector (BSED) for backscattered electrons (BSE)(Duarte et al., 2026).

### DNA extraction and phylogenetic analysis

The cyanobacterial cells were harvested from the culture, and total genomic DNA (gDNA) of the strain was extracted using the PureLink Genomic DNA kit (Invitrogen, Waltham, Massachusetts, USA), following the manufacturer’s instructions for Gram-negative bacteria. Specific cyanobacterial primers were used for gene amplification, specifically primers 27SF (Neilan et al., 1997) and 23SR (Taton et al., 2003). The PCR reaction was performed with the following conditions: initial denaturation at 94°C for 5 min, followed by 10 cycles of denaturation at 94°C for 45s, annealing at 57°C for 45s and extension at 72°C for 2 min. This was followed by an additional 25 cycles of denaturation at 92°C for 45s, annealing at 54°C for 45s, and extension at 72°C for 2 min, with a final extension step at 72°C for 7 min. PCR product was submitted to electrophoresis (80 V, 1 hour), followed by gel excision. Fragment of the expected size were excised from the gel and purified using the NZYGelpure kit (Nzytech, Portugal), following the manufacturer’s instructions. Finally, the purified fragment was sent for sequencing (separately) with primers 359 F and 781 R (Nübel et al., 1997), 1494 R and 27SF (Neilan et al., 1997), 23SR (Taton et al., 2003), and 1114 F (Lane, 1991). The sequencing was performed by Sanger dideoxy sequencing at GATC Biotech (Ebersberg, Germany), and the nucleotide sequence obtained was manually inspected for quality and assembled using Geneious Prime 2023.2.1 software (Biomatters Ltd, Auckland, New Zealand) (Hentschke et al., 2025). The obtained sequence was deposited in GenBank (National Center for Biotechnology Information, NCBI).

The phylogenetic analyses were conducted in two rounds. In the first round, we aimed to position LEGE 10371 among the cyanobacterial genera. For that, we aligned the 16S rRNA gene sequence of LEGE 10371 with sequences of cyanobacterial reference strains and additional sequences retrieved from GenBank (NCBI) by BLAST. This alignment contained 440 sequences. The phylogenetic reconstruction was performed using the FastTree method (Price et al., 2009), with the bootstrap value set to the default of 1000 as per the manual. The resulting tree was edited using iTOL(Letunic & Bork, 2021). In the second round of analysis, to obtain confirmation of the taxonomic status of the isolates, we aligned their sequences with Pseudanabaenales genera. This alignment contained 45 sequences. Then, the phylogenetic trees were built using Maximum Likelihood (ML) and Bayesian Inference (BI) analysis. GTR+G+I evolutionary model was selected by MEGA11: Molecular Evolutionary Genetics Analysis version 11 (Tamura et al., 2021), according to the AIC criterium. The robustness of ML tree was estimated by bootstrap percentages, using 1000 replications using IQ-Tree online version v1.6.12 (Trifinopoulos et al., 2016). The BI tree was constructed in two independent runs, with four chains each, for 5×10^6^ generations, burnin fraction set to 0.25, sample frequency 1000, using MrBayes (Ronquist et al., 2012) in Cipres Gateway(Miller et al., 2010). The processing and visualization of these trees were made using FigTree v1.4.4 (http://tree.bio.ed.ac.uk/software/figtree).

For all these analyses, the sequences were aligned using MAFFT(Katoh et al., 2002) and the outgroup used was *Gloeobacter violaceus* PCC 8105 (AF132791). The alignments were performed with the full-length 16S rRNA gene sequences of the strains in the phylogenetic trees. The 16S rRNA gene identities (p-distance) were calculated using MEGA11. The 16S-23S rRNA ITS secondary structures were folded using MFOLD(Zuker, 2003), according to Lukesova et al. (2009).

### Cyanotoxin-producing genes detection

In order to determine the safety of the studied strain and its potential for biotechnological applications, we used PCR amplification and 1.5% agarose gel electrophoresis (100 V, 20 min) to detect cyanotoxin-producing genes by identifying specific DNA bands (Benredjem et al., 2023). For cylindrospermopsin, the amplification of a cyrJ fragment (584 bp) was performed using the primers cynsulF and cylnamR (Mihali et al., 2008). PCR validation used *Cylindrospermopsis raciborskii* strain LEGE 97047 as positive control. For saxitoxin, the amplification of a sxtI fragment (200 pb) was performed using the primers SxtI682F and (Lopes et al., 2012).PCR validation was obtained using *Aphanizomenon gracile* strain LMECYA 040 as positive control. For anatoxin, the amplification of an anaC fragment (366 bp) was performed using the primers anaC-genF and anaC-genR (Rantala-Ylinen et al., 2011), with *Anabaena* sp. strain LEGE X-002 as positive control for PCR. For microcystin the amplification of a mcyA fragment (472 pb) was performed using the primers HEPF and HEPR (Jungblut & Neilan, 2006). PCR validation was obtained using the strain *Microcystis aeruginosa* LEGE 00063 as positive control (Hentschke et al., 2025).

### MALDI-TOF Mass Spectrometry

Two aliquots of cyanobacterial culture were transferred into two 2 mL Eppendorf tubes and centrifuged at 13,000 g for 2 min. The supernatant was discarded, and the cell pellets (∼0.5 mL of wet biomass) were washed with ultrapure water, followed by centrifugation under the same conditions. After removal of the supernatant, 900 µL of acetone was added to each tube, and the samples were mixed using a mechanical shaker. The suspensions were centrifuged at 12,000 g for 10 min, and the supernatants were discarded. Subsequently, 1,5 mL of acetone was added, and the pellets were resuspended by mechanical shaking and centrifuged again at 12,000 g for 10 min. The supernatants were discarded, and the pellets were allowed to dry completely. The dried extracts were then resuspended in 30 µL of 70% formic acid and incubated for 5 min, followed by the addition of 30 µL of acetonitrile. The samples were centrifuged at 13,000 g for 2 min, and the final supernatants were collected.

For each tube (2 tubes), one microliter of the sample was deposited on the MALDI plate in duplicate (n=4), dried using MBT FAST™ Shuttle benchtop device, covered with α-hydroxycinnamic acid (HCCA) matrix, and allowed to dry again.

MALDI-TOF MS of cyanobacterial cells was performed on a MALDI Biotyper® Sirius instrument (Bruker Daltonics, Bremen, Germany). Mass Spectra were acquired in a positive ion mode across a mass/charge ratio (m/z) of 2,000 to 20,000 Da and the instrument was calibrated externally using the Bacterial Test Standard covering average masses between 3637.8 and 16952.3 Da (Bruker Daltonics, Bremen, Germany). The acquisition software was flexControl 3.4 and the spectral data were checked and saved using flexAnalysis 3.4 (all by Bruker Daltonics).

#### Biomass growing and targeted extraction

For biomass growing, a culture volume of 4 L was prepared from the 40 mL inoculum, with continuous aeration for 4 weeks. Cultures were kept under controlled temperature conditions (21 °C ± 2 °C), with photoperiod of 16:8 h (light/dark cycle) and light intensity of 20 µmol photons m−2 s−1 (AsenseTek Lighting Passport, Biosystems, United Kingdom) with fluorescent light (Acardia, United Kingdom). The biomass was harvested by centrifugation at 4700 rpm for 10 min at 4 °C (Sorvall BIOS 16 Bioprocessing Centrifuge, Thermo Fisher Scientific, Bremen, Germany). A washing step with deionized water was performed to remove the salt. The fresh biomass was dried in a freeze dryer (Lyoquest-55, Telstar Technologies, S. L., Spain) and stored at −20 °C prior to extracts preparation.

#### Acetone fraction

Acetone extraction was performed using 2 g of dry biomass, which was immersed in 40 mL acetone (99.5%, Panreac, Barcelona, Spain) and subjected to ultrasound-assisted extraction with a MS-72 probe (Bandelin Sonopuls, Berlin, Germany) with an amplitude of 80% and no pulse, for 5 min. The sample was kept on ice throughout the procedure to prevent temperature increases caused by sonication, thereby preserving thermolabile compounds and minimizing potential oxidative degradation. Subsequently, centrifugation (10000 rpm, 5 min, 4 °C) was employed to remove cellular debris, using a Thermo Scientific™ HERAUS Megafuge™ 16R centrifuge (Waltham, MA, USA). The supernatant was filtered through a 0.22 µm pore membrane (FilterBio, Nantong, China) and reserved. The biomass was re-extracted three times; filtrates were combined and evaporated under reduced pressure using a BUCHI R-210 Rotary Evaporator (Cambridge, MA, USA) set at 200 mbar, with the water bath at 30 °C and the condenser at −8 °C, to remove the solvent while preserving the integrity of the extracted compounds. The resulting residue was weighed and stored at −20 °C until chemical and biological analyses.

#### Water fraction

Following acetone extraction, the residual biomass was left in a fume hood overnight to evaporate the remaining solvent. The dry pellet was then extracted with 40 mL of ultrapure water following the same procedure; this was repeated three additional times. After extraction, the mixture was centrifuged, and the supernatants collected, frozen and lyophilized. Dried extracts were weighed and stored until analysis.

### Cytotoxicity to Human Keratinocytes (HaCaT)

To evaluate the safety profile of the acetone and aqueous cyanobacterial fractions, an in *vitro* cytotoxicity screening was conducted with human keratinocytes (HaCaT, ATCC).

### Cells Culturing

Cell cultures were maintained in Dulbecco’s Modified Eagle Medium with high glucose (DMEM; Gibco), supplemented with 10% fetal bovine serum (FBS; Gibco), 1% penicillin–streptomycin (final concentrations of 100 IU mL⁻¹ and 10 mg mL⁻¹, respectively; Gibco), and 0.1% amphotericin B (Gibco). Cells were incubated at 37 °C in a humidified environment containing 5% CO₂, with medium replacement every 48 h. Once cultures reached approximately 80–90% confluence, cells were dissociated using TrypLE Express Enzyme (Gibco), pelleted by centrifugation at 1200 rpm for 5 minutes. (Eppendorf 5430), resuspended in fresh DMEM, and either reseeded or transferred to new culture flasks. All manipulations were conducted aseptically in a Thermo Scientific™ Safe 2020 laminar flow cabinet (Thermo Fisher Scientific, Waltham, MA, USA).

### Cytotoxicity Assessment by the MTT Assay

Cells viability was determined using the MTT colorimetric assay, based on the metabolic reduction of 3-(4,5-dimethylthiazol-2-yl)-2,5-diphenyltetrazolium bromide (MTT, 98%; Thermo Scientific), following previously established protocols (Morone et al., 2022). Cells were seeded in 96-well plates at a density of 2.5 × 10⁴ cells mL⁻¹ and allowed to attach for 24 h.The culture medium was then replaced and cells were exposed to five serial concentrations (12.5–200 µg mL⁻¹) of each fraction for 24 and 48 h. The acetone fraction was resuspended in dimethyl sulfoxide (DMSO; Gibco) and diluted in DMEM. Working solutions of the acetone fraction were prepared from a 20 mg mL⁻¹ stock solution, whereas those of the aqueous fraction were prepared from a 10 mg mL⁻¹ stock in PBS. Control conditions included DMEM and 1% DMSO in DMEM, while 20% DMSO in DMEM was used as a positive control for cytotoxicity.

At the end of the exposure period, 20 µL of MTT solution (1 mg mL⁻¹ in PBS) were added to each well, followed by incubation for 3 h. The supernatant was removed, and the resulting formazan crystals were dissolved in 100 µL of DMSO. Absorbance was measured at 550 nm using a Synergy HT multi-detection microplate reader (BioTek, Bad Friedrichshall, Germany) equipped with Gen5™ software (version 2.0). All experiments were performed in quadruplicate, and data were reported as mean ± standard deviation. Cells viability was calculated as the percentage relative to the corresponding solvent control.

### Chemical characterization

#### Carotenoid and Chlorophylls Profiling by HPLC-PDA

The pigment composition of acetone extracts obtained from cyanobacteria was analyzed by high-performance liquid chromatography (HPLC) using a Waters Alliance 2695 system coupled to a photodiode array (PDA) detector (Waters, USA), in accordance with the procedure described by (Morone et al., 2022). Chromatographic data were acquired and processed with Empower^®^ 2 software (Waters, NJ, USA), and UV–Vis spectra were recorded for all detected peaks over a wavelength range of 250–750 nm. Pigments were tentatively assigned based on comparisons of retention times and absorption spectra with those of commercially available reference standards. Quantification of carotenoids was performed by monitoring absorbance at 450 nm and comparing peak areas with those obtained from external standards.

Authentic standards of lutein, chlorophyll-*a*, zeaxanthin, β-cryptoxanthin, echinenone, and β-carotene (Extrasynthese, Genay, France; Sigma-Aldrich, St. Louis, MO, USA; DHI, Hørsholm, Denmark) were used for compounds identification and quantification. Unidentified xanthophylls were quantified as zeaxanthin, selected as the predominant xanthophyll, while chlorophyll-*a* derivatives were quantified as chlorophyll-*a*, the principal chlorophyll present in cyanobacteria. Calibration curves were generated using five standards concentrations covering the expected range in the samples. The resulting calibration curves and coefficients of determination (*r*²) for the analyzed pigments are presented in Table 1, where *x* corresponds to analyte concentration and *y* to chromatographic peak area.

**Table 1.** Calibration curves of authentic standards used for quantification of different carotenoids and chlorophylls.

| Standards | Calibration Curve | $r^2$ |
| --- | --- | --- |
| **Lutein** | $y = 83327993.12x + 1378.83$ | 0.9998 |
| **β-Carotene** | $y = 54734949.60x - 58044.75$ | 0.9995 |
| **Chlorophyll-*a*** | $y = 6071813.05x + 19901.47$ | 0.9971 |
| **Echinenone** | $y = 75013851.88x - 31327.01$ | 0.9962 |
| **Zeaxanthin** | $y = 86507689.89x - 51378.29$ | 0.9996 |

#### Phycobiliproteins and Chlorophylls Profiling by Spectrophotometry

Due to their water-soluble nature, phycobiliproteins (PBPs) were quantified spectrophotometrically in the aqueous extracts. Samples were diluted with distilled water to obtain a final concentration of 0.5 mg mL⁻¹. Absorbance readings were recorded at 562, 615, 652 and 675 nm using cuvettes with a 1 cm path length. Measurements were carried out in duplicate, and PBP concentrations were calculated according to previously established equations of Bennett & Bogobad, (1973) subsequently adjusted by Lauceri and co-workers (Lauceri et al., 2018) to reduce the interference of chlorophyll-a in PC and APC determination:

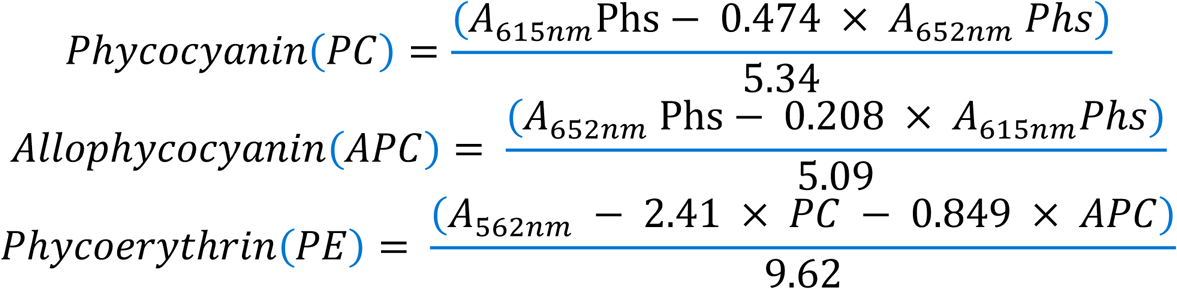

Where:

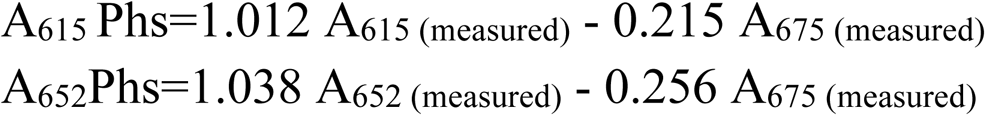

For Chrolophylls, quantification was based in the universal equations suggested by Ritchie, 2008:

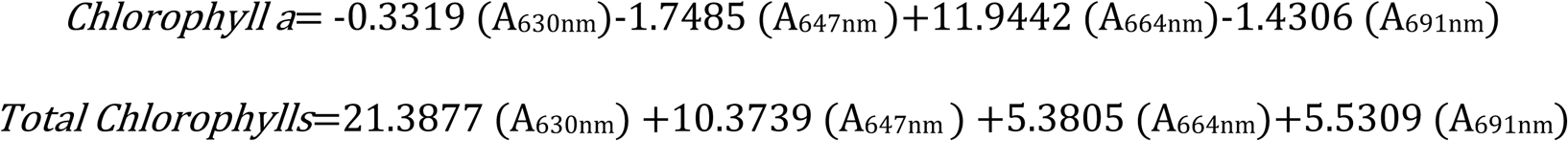

Results were expressed as micrograms of the corresponding phycobiliprotein or chlorophylls per milligram of dry extract.

#### Biological Activities

The ability of cyanobacteria fractions to counteract biologically significant reactive species and enzymes involved in the inflammatory framework was assessed by testing their capacity to scavenge *in vitro*–generated superoxide anion (O_₂_^•⁻^) and nitric oxide (^•^NO) radicals.

#### O_2_^−^ Scavenging

Superoxide anion radical scavenging activity was evaluated according to a previously reported protocol (Morone et al., 2022). Aqueous fractions were resuspended in water, whereas acetone ones were solubilized in DMSO. Serial dilutions of each fraction were prepared in phosphate buffer (19 µM, pH 7.4). Scavenging activity was determined by measuring the inhibition of nitro blue tetrazolium bromide (NBT) reduction induced by O_₂_^•⁻^, using a Synergy HT multi-detection microplate reader (BioTek, Bad Friedrichshall, Germany) operated with Gen5™ software. Kinetic measurements were performed at room temperature for 2 min at 562 nm. Gallic acid was used as the reference antioxidant. Results were expressed as the percentage of O_₂_^•⁻^neutralization relative to the untreated control, and dose–response curves were generated using GraphPad Prism® software (version 10.3.1 for MacOs).

#### • NO Scavenging

Nitric oxide radical generated from sodium nitroprusside (SNP) was quantified using the procedure reported by Lopes et al. (2012). Aqueous fractions were resuspended in water, whereas acetone ones were solubilized in DMSO. Serial dilutions were prepared in phosphate buffer (100 mM, pH 7.4). After the incubation period, the absorbance of the reaction-derived chromophore was determined using the Griess reaction at 562 nm with a Synergy HT multi-detection microplate reader (BioTek, Bad Friedrichshall, Germany) operated using Gen5™ software. Blanks were included for each tested concentration. Radical scavenging activity was expressed as the percentage reduction relative to the untreated control, with quercetin employed as the reference antioxidant.

#### LOX inhibition

The inhibitory effect of cyanobacteria fractions in LOX followed a procedure previously described by Fernandes and co-workers (Fernandes et al., 2018). Serial dilutions of the fractions under study were pre-incubated in a reaction mixture containing 20 µL of fraction, 200 µL of phosphate buffer (pH 9.0) and 20 µL of soybean lipoxygenase (150 U/20 µL), for 5 min at room temperature. After the incubation period, 20 µL of substrate (linoleic acid, 4.18 mM in absolute ethanol) was added to start the reaction. Absorbance was measured continuously at 234 nm with a Synergy™ HT plate reader (Biotek Instruments; Winooski, VT, USA) operated by Gen5 Software (version 2.0) over 3 min. LOX inhibition was calculated by comparing the rate of reaction in the presence of the fractions relative to the untreated control. Quercetin was used as positive control.

## Results

### Taxonomic Results

The 16S rRNA gene first-round phylogenetic tree confirmed that *Pseudanabaena*, *Tumidithrix*, *Thalassoporum, Portococcus* and *Pseudanabaenococcus* are phylogenetically distinct. Particularly, the *Thalassoporum* clade is positioned at the base of a cluster containing the other genera. Within the *Thalassoporum* clade (ML=100), the strain LEGE 10371 was clearly separated from the type species *Th. komareki* TAUMAC 1515 and from *Th. mexicanum* PCC 7367 (Fig. S1).

The second-round ML and BI phylogenetic analyses were in agreement with the previous results. In both trees, *Pseudanabaena*, *Tumidithrix*, *Thalassoporum, Portococcus* and *Pseudanabaenococcus* formed distinct and strongly supported clades, confirming their status as separate genera. Strain LEGE 10371 was placed again within the *Thalassoporum* clade (ML= 100%, BI = 1) but clearly separated from the other described species of the genus, indicating that it represents a novel species (Fig. 1).

**Figure 1.**
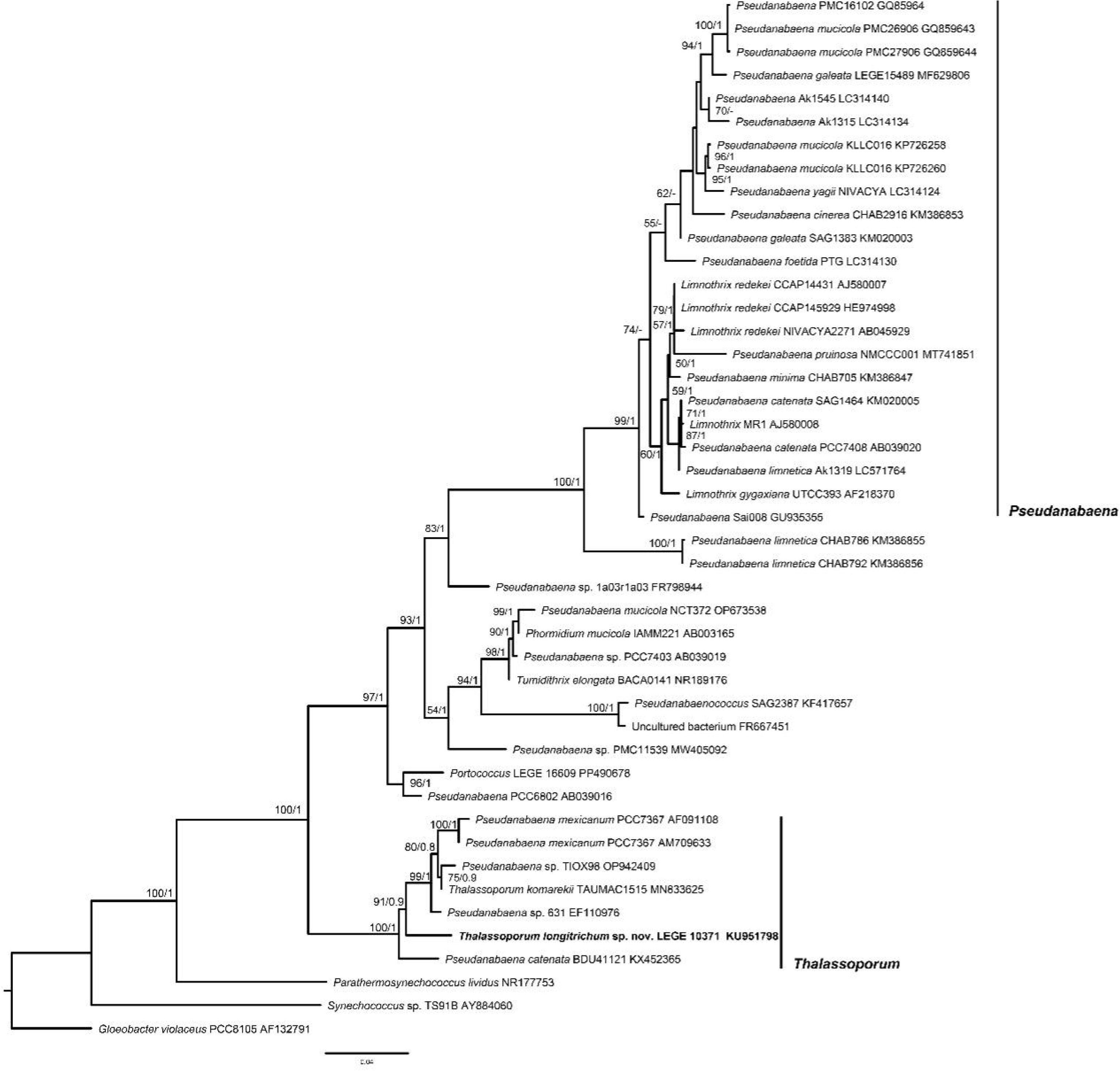
16S rRNA gene phylogenetic tree with 41 sequences. Bootstrap values and BI posterior probabilities are presented at the nodes.

The 16S rRNA gene identity analysis confirmed that LEGE 10371 is distinct, exhibiting low sequence identity with other *Thalassoporum* species, with a maximum value of 97.2% to *Th. komarekii* (Table S1). The dissimilarity values of the 16S–23S ITS sequences of the three reference strains could not calculated. The sequences are so divergent that they could not be reliably aligned, also indicating that they represent three distinct species.

This divergence in the 16S–23S ITS region was reflected in the distinct predicted secondary structures between *Thalssoporum* species. In the D1–D1′ helix, the unilateral basal bulge (UBB) was not conserved among the *Thalassoporum* sequences compared. For example, the UBB region of strain LEGE 10371 contains a short sequence (5′-ACUCC-3′) without a complementary opposing nucleotide. In contrast, the type species *Th. komarekii* shows an insertion of a cytosine in this region (5′-CACUCC-3′), with the UBB opposed by an adenine. In *Th. mexicanum*, the inserted cytosine is replaced by an adenine, resulting in the sequence 5′-AACUCC-3′, which is likewise opposed by an adenine at the 5′ side of the molecule. Additional differences among strains were observed in other regions of the D1–D1′ helix; notably, the terminal loop was unique for each species. Moreover, *Th. komarekii* was the only species exhibiting a loop in the medial region of the helix. The Box B and V3 helices were also highly variable, showing differences in sequence, length and structure among strains (Fig. 2). Altogether, these structural and sequence divergences strongly support the separation of the analyzed strains into distinct species.

**Figure 2.**
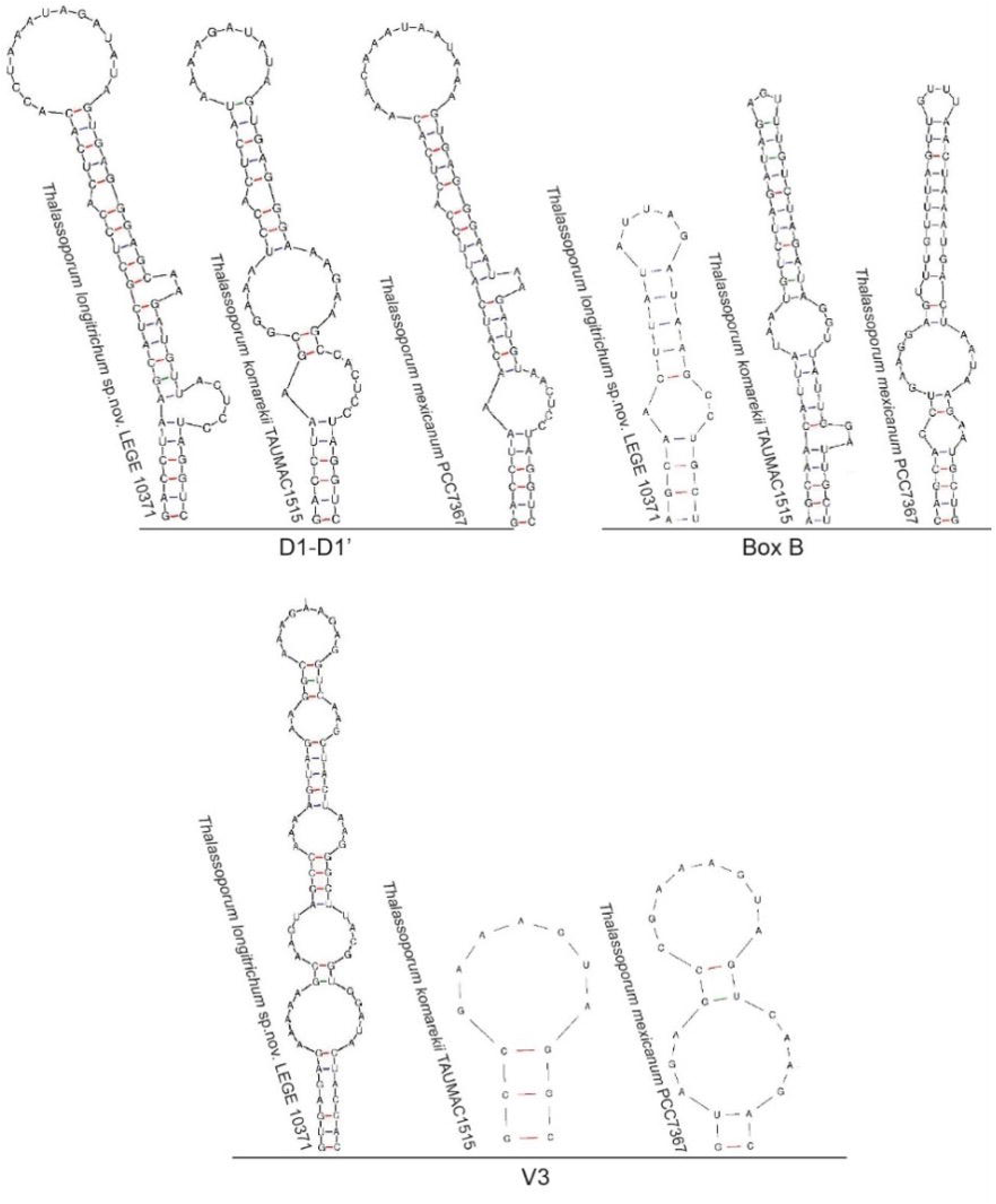
16S-23S ITS secondary structures of *Thalassoporum* species.

**Figure 3.**
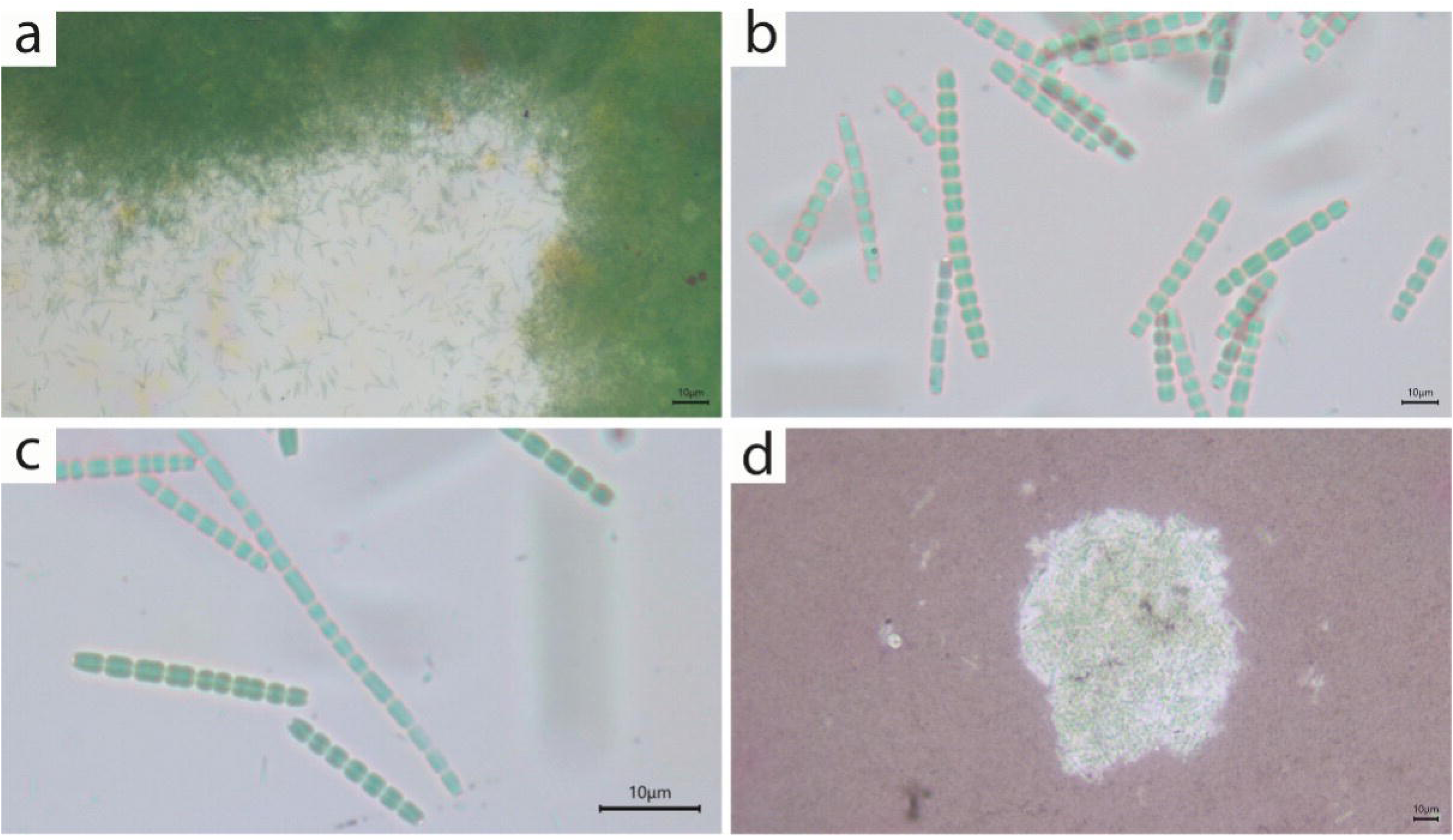
Morphology of *Th. longitrichum* sp. nov. LEGE 10371 in optic microscope. a. General view of colony. b,c. Details of short (<10 cells) and long (>10 cells) trichomes. d. Slide preparation with China Ink showing a thin layer of diffluent mucilage surrounding the colony.

**Figure 4.**
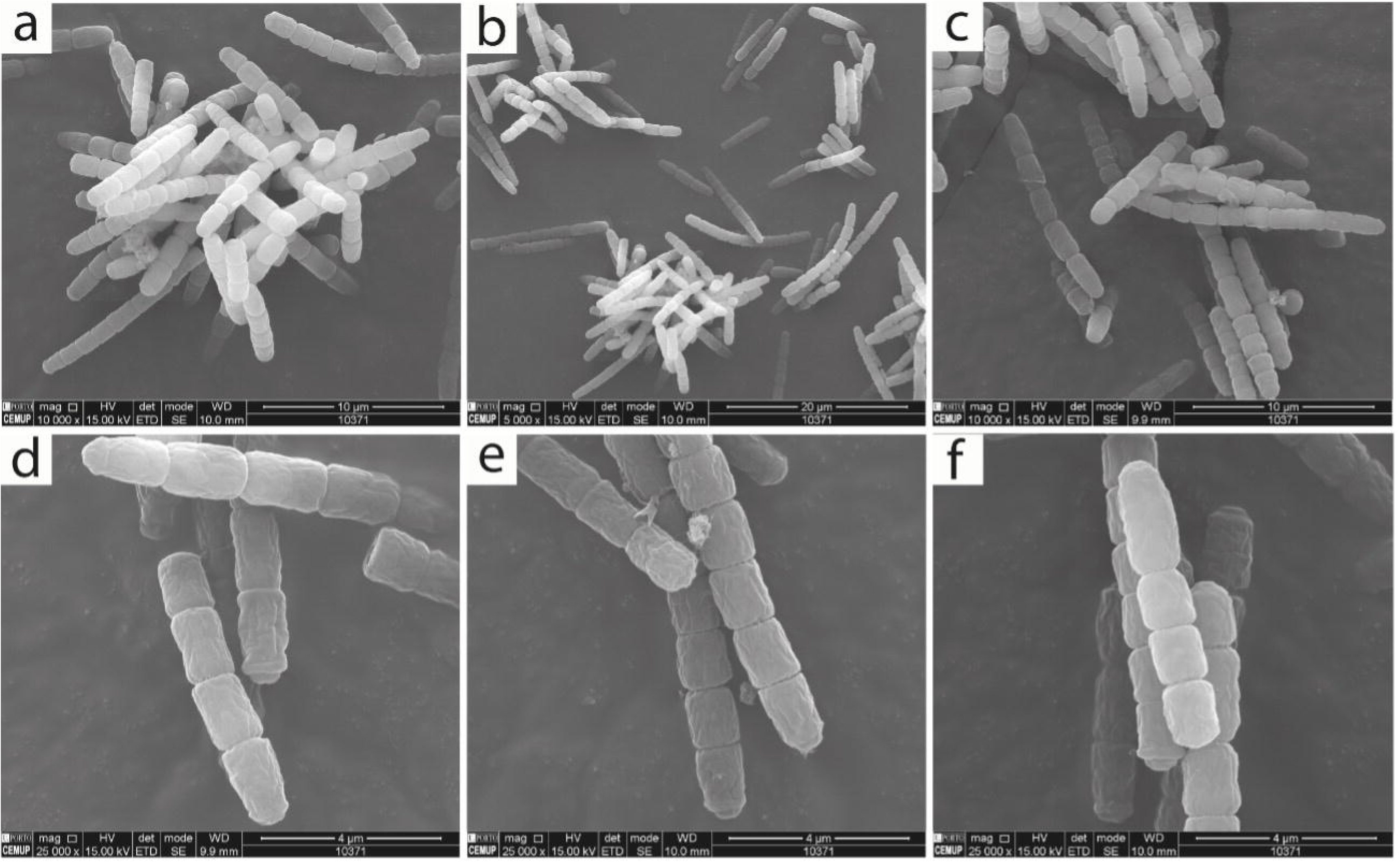
Morphology of *Th. longitrichum* sp. nov. LEGE 10371 in SEM. a-c. Colonies with short filaments. d,e. Filaments with conical apical cells. f. detail of trichomes.

The general morphology of LEGE 10371 conforms to the diagnostic traits of the genus *Thalassoporum*, including homocytic trichomes, strong constrictions and absence of a firm sheath. However, LEGE 10371 differs from *Th. mexicanum* and *Th. komarekii* by the absence of aerotopes, which are present in both latter species. Additionally, the trichomes of LEGE 10371 are longer (>10 cells), whereas trichomes in *Th. mexicanum* and *Th. komarekii* comprise up to 10 cells. (Table 2). Reproduction occurs through trichomes fragmentation, but hormogonia were not observed. Ecologically, LEGE 10371 represents the only epizoic strain reported for the genus to date, being isolated from the surface of a marine sponge.

**Table 2.** Morphological and habitat comparisons between *Thalassoporum* species.

| Character | *Th. longitrichum* sp. nov. LEGE 10371 | *Th. mexicanum* PCC 7367 | *Th. komarekii* TAU-MAC 1515 (type of the genus) |
| --- | --- | --- | --- |
| **Trichome type** | Uniseriate, homocytic; straight to slightly curved; composed of long series of cells (>10). No attenuation at ends. | Homocytic, uniseriate; straight; 3–10 cells per trichome; not attenuated. | Homocytic, uniseriate; 3–10 cells; not attenuated. |
| **Trichome width** | 1.3–1.5 μm | 1.3–1.9 μm | 1.08–1.74 μm |
| **Sheath** | Absent around individual trichomes | Absent or very thin | Absent |
| **Cell shape** | Barrel-shaped to slightly elongated, cylindrical; longer than wide. | Barrel-shaped to cylindrical; rounded ends | Cylindrical; rounded ends |
| **Cell length** | 2–3 μm | 1.9–3.7 μm | 1.3–3.24 μm |
| **Constrictions at cross-walls** | Deeply constricted | Deeply constricted | Deeply constricted |
| **Apical cell** | Slightly rounded | Flat rounded or conical-rounded | Flat rounded or conical-rounded |
| **Gas vesicles** | Not present | Present (polar) | Present (polar) |
| **Trichome flexibility** | Straight to slightly curved; moderately rigid; not coiled | Moderately flexible | Moderately flexible |
| **Thylakoids** | Not observed | 6–10 peripheral | 5–6 peripheral |
| **Reproduction** | Fragmentation | Fragmentation & hormogonia | Not reported |
| **Mucilage** | Diffuse colonial mucilage | Weak/occasional | Absent |
| **Colony appearance** | Dispersed trichomes forming loose clusters; no dense mats | Sparse free filaments; no mucilage; not forming compact colonies | Loosely arranged free filaments; no sheath; no mucilage |
| **Habitat (reported)** | On marine sponge, intertidal zone | Marine, intertidal zone | Marine, rocky sublittoral |
| **Locality** | Praia da Memória, Portugal | Puerto Peñasco, Mexico | Porto Valitsa, Greece |

The four MALDI-TOF MS spectra obtained exhibited highly consistent protein profiles, consistently displaying high-intensity peaks at approximately 3,000, 4,000, 6,000 and 8,000 m/z (Fig. S2). These peaks represent strong candidates for diagnostic markers of strain LEGE 10731 and may potentially distinguish it from other species within the genus. To confirm their taxonomic specificity, comparative spectra from *Th. komarekii* and *Th. mexicanum* should be obtained and analyzed in the future.

Based on the polyphasic approach comprising molecular, morphological and habitat comparisons, it was evident that LEGE 10371 is a novel species of *Thalassoporum*.

## Description

### Order Pseudanabaenales

#### Family Thalassoporaceae

*Thalassoporum longitrichum* sp. nov. G. S. Hentschke & C. Tenorio Rodas **Figures 3 and 4**

##### Diagnosis

This species differs from the others of *Thalassoporum* by the longer trichomes (>10 cells) and the lack of aerotopes.

##### Description

Colonies embedded in a narrow diffuse mucilaginous matrix, formed by irregularly disposed trichomes, sometimes in compact aggregates. Trichomes uniseriate, straight to slightly curved, composed of short (<10) or long series of cells (>10), not attenuated at the ends and lacking firm sheaths. Cells constricted at the cross-walls, forming narrow transverse septa that sharply delimit adjacent cells. Cell barrel-shaped to slightly elongated (cylindrical), usually longer than wide, rarely shorter than wide, with parallel lateral walls. Apical cells with rounded ends, sometimes conical. Cell dimensions: 1.5–3 µm long; 1.3–2 µm wide. Cell content homogenous; no gas vesicles observed. Reproduction by trichome fragmentation.

##### Etymology

The specific epithet *longitrichum* is derived from the Latin *longus* (long) and the Greek *thrix* (hair), referring to the long trichomes observed in this species.

##### Holotype

Collected from the surface of a marine sponge at Praia da Memória, Portugal (41.23119 N 8.721750 W) in 2010 by Ana Regueiras. Deposited in the University of Porto herbarium in a metabolically inactive state (lyophilized) under the code PO-T5292.

##### Reference strain

LEGE 10371; KU951798 (16S rRNA gene)

Habitat: On marine sponge, intertidal zone.

Beyond the description of the new species, phylogenetic and 16S rRNA gene identity analyses confirmed that *Pseudanabaena* and *Limnothrix* are not distinguishable as separate genera. Notably, strains of the type species *L. redekei* (CCAP 1443/1, CCAP 1459/29, and NIVA-CYA 227/1) consistently clustered within the *Pseudanabaena* clade with strong phylogenetic support (ML = 100; BI = 1) (Figs. 1 and S1). Moreover, 16S rRNA gene sequence identity between *Limnothrix* and *Pseudanabaena* strains was very high, ranging from 96.1% to 99.9% (mean = 97.7%) (Table S2).

The 16S–23S ITS secondary structures of *Pseudanabaena/Limnothrix* clade were very similar and a common diagnostic marker was identified in the D1–D1′ helices. In the UBB region, all *Pseudanabaena* and *Limnothrix* strains shared the sequence 5′-CAUCCA-3′, with no nucleotide opposing it on the 5′ side of the molecule. The only exceptions were strains CHAB786 and CHAB792, which exhibited a deletion of the “U”, resulting in the sequence 5′-CACCA-3′ (Fig. 5). The other helices exhibited higher variability, a pattern also observed in the secondary structures of the different *Thalassoporum* species (Fig. 6, 7).

**Figure 5.**
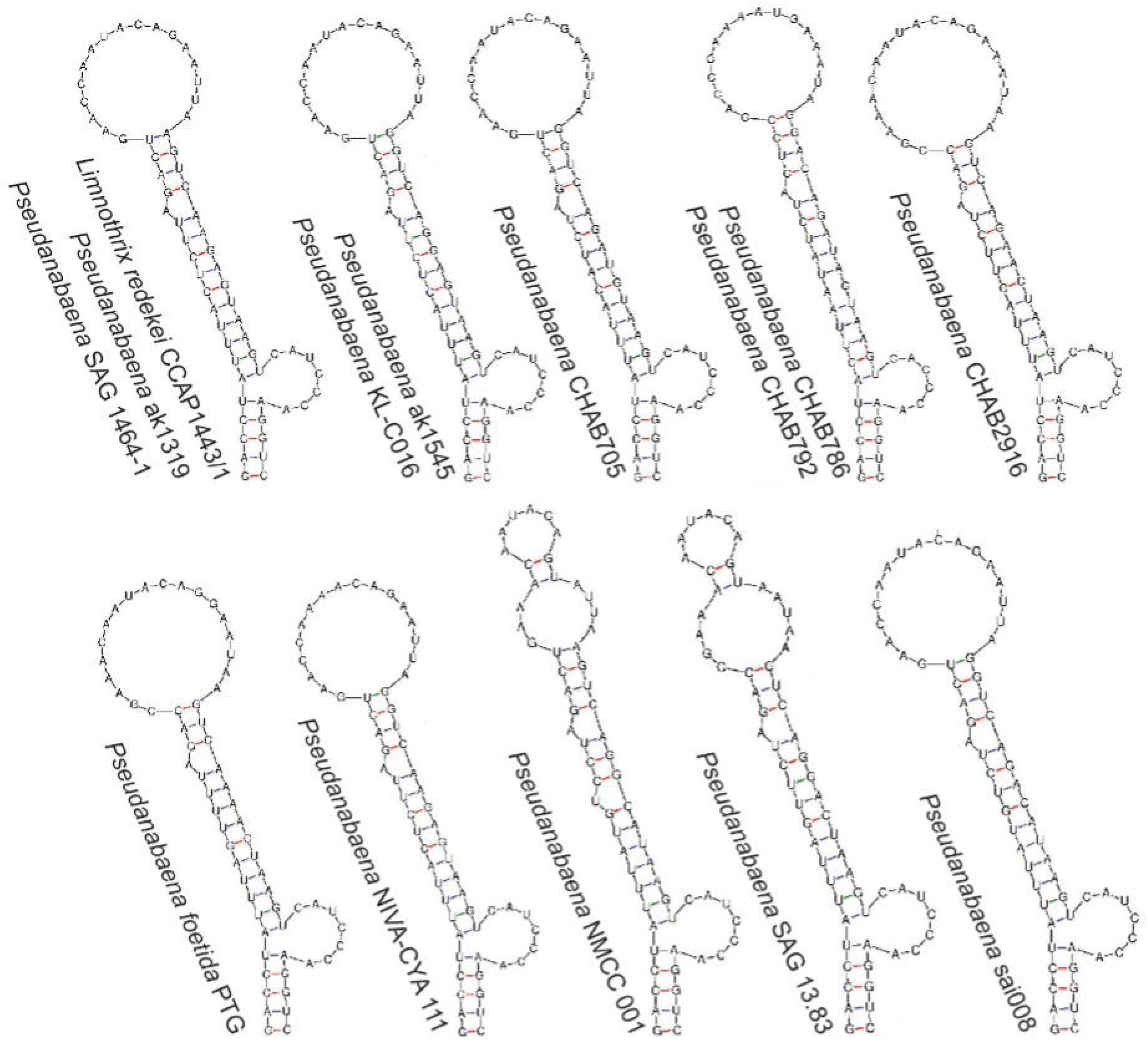
D1-D1’ helices of 16S-23S ITS secondary structures of *Pseudanabaena* and *Limnothrix*.

**Figure 6.**
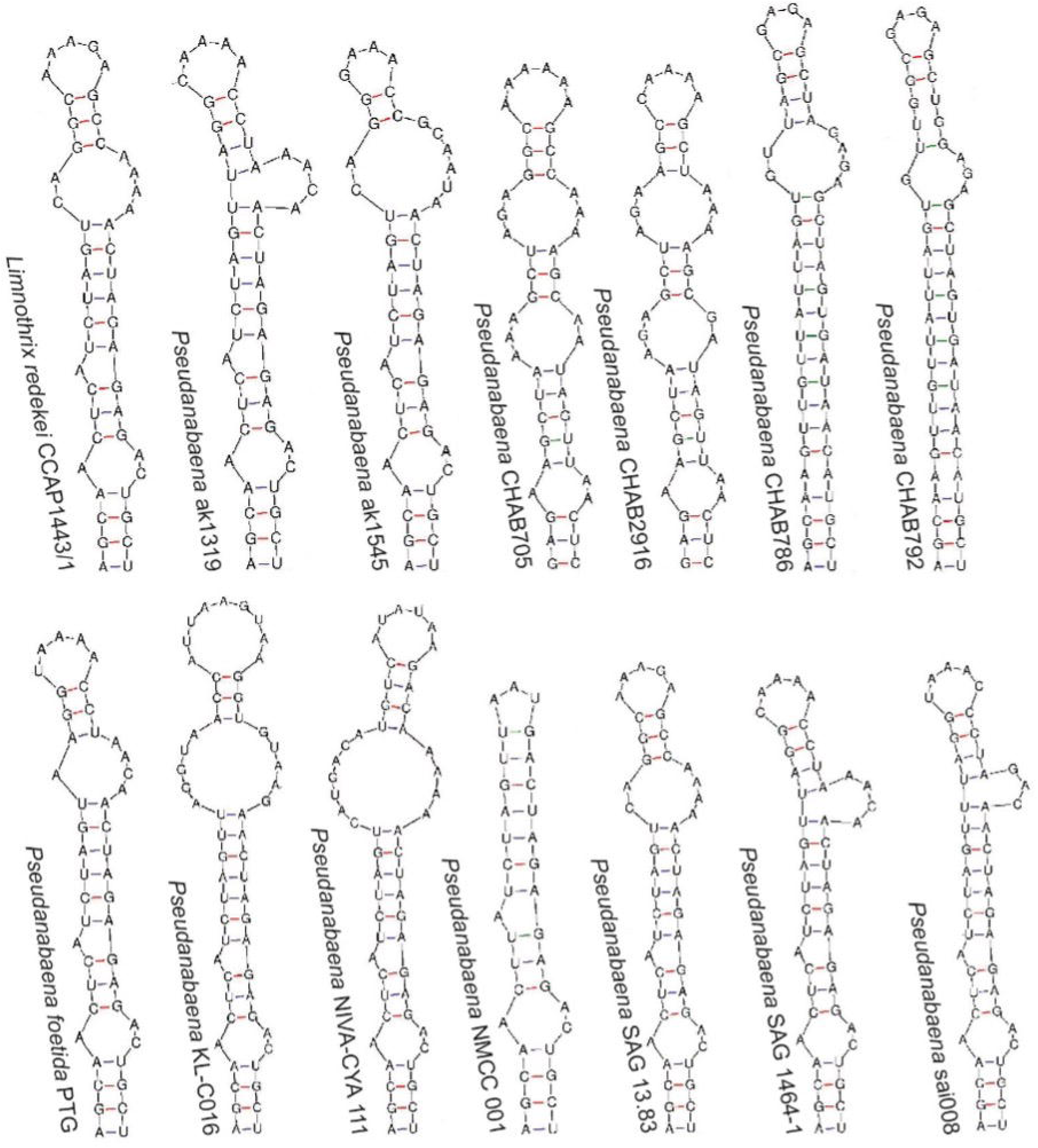
Box B helices of 16S-23S ITS secondary structures of *Pseudanabaena* and *Limnothrix*.

**Figure 7.**
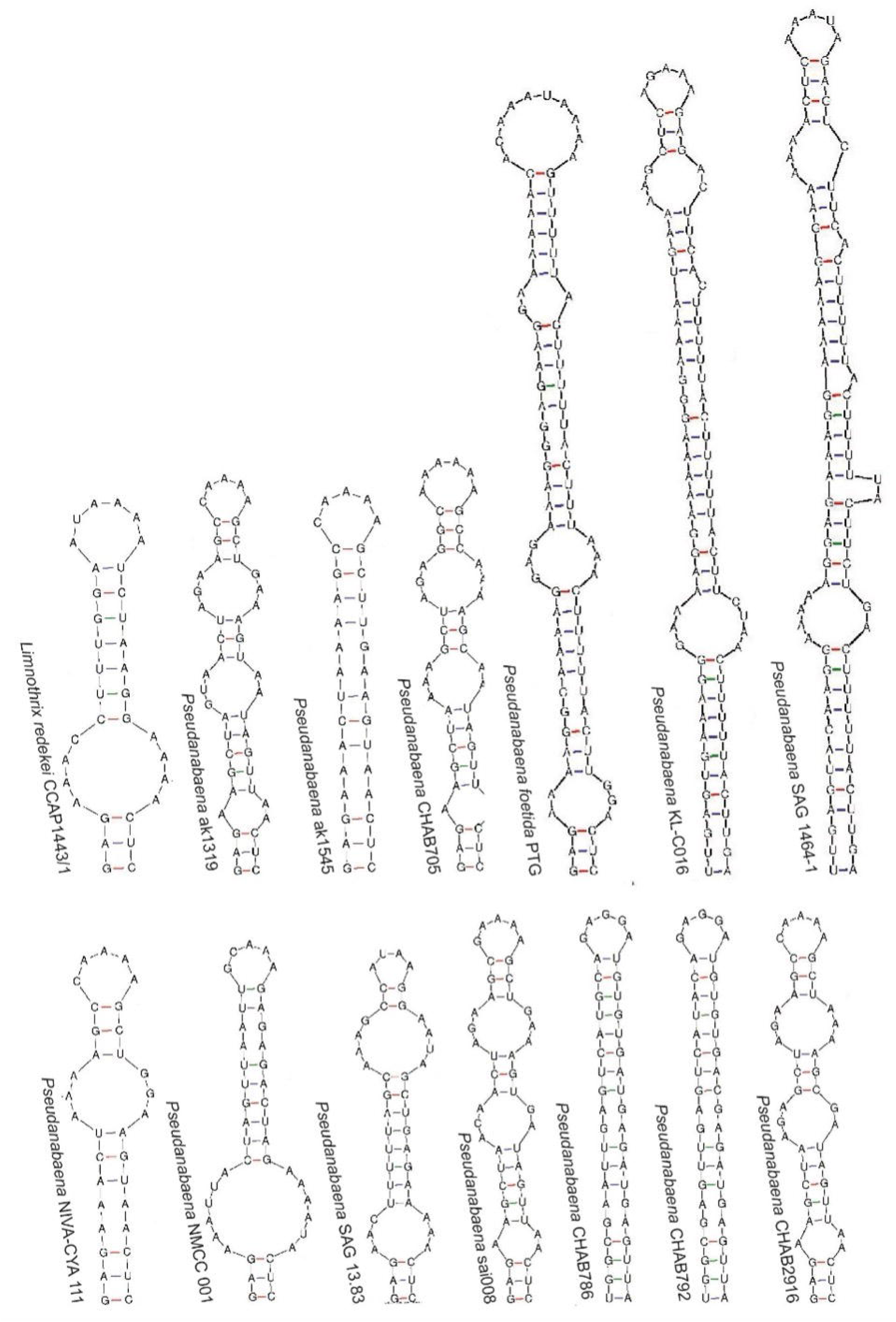
V2 helices of 16S-23S ITS secondary structures of *Pseudanabaena* and *Limnothrix*.

Given the molecular and morphological identity between *Pseudanabaena* and *Limnothrix*, *Limnothrix* cannot be recognized as a distinct genus and is herein transferred to *Pseudanabaena*. Consequently, we present an emended description for *Pseudanabaena* with addition of the *Limnothrix* characters.

### New combinations

***Pseudanabaena*** Lauterborn, 1915

Published in Lauterborn, R. (1915). Die sapropelische Lebewelth. Ein Beitrag zur Biologie des Faulschlammes natürlicher Gewässer. Verhandlungen des Naturhistorisch-medizinischen Vereins zu Heidelberg, Neue Folge 13: 395-481, plate 3.

### Synonym

*Limnothrix* M.-E.Meffert, 1987. Published in: Meffert, M.-E. (1987). Planktic unsheathed filaments (Cyanophyceae) with polar and central gas vacuoles. I. Their morphology and taxonomy. Archiv für Hydrobiologie Supplement 76: 315–346.

***Pseudanabaena redekei*** (Meffert) G. S. Hentschke & C. Tenorio Rodas **comb. nov.** Basionym: *Limnothrix redekei* (Goor) Meffert 1987: 327, figs 38–62; pl. 4

Published in: Meffert, M.-E. (1987). Planktic unsheathed filaments (Cyanophyceae) with polar and central gas vacuoles. I. Their morphology and taxonomy. Archiv für Hydrobiologie Supplement 76: 315–346.

### Emmended description for *Pseudanabaena*

Filaments solitary or forming aggregates or mats, straight to slightly curved, less frequently waved, unbranched, cylindrical. Trichomes usually short consisting of very few cells, or long with many cells. Cross-walls variably constricted, being clearly constricted in some taxa and weakly constricted or unconstricted in others. Sheaths absent or thin, colorless, or indistinct, sometimes not discernible under light microscopy. Sometimes with diffluent mucilage. Cells cylindrical to barrel-shaped cells, usually longer than wide, rarely tending to isodiametric, with rounded ends. Cell width up to 3.5 µm. Apical cells morphologically similar to intercalary cells, without calyptra or terminal differentiation. Ends rounded or flattened, rarely conical. Reproduction occurs by fragmentation of trichomes into hormogonia. Aerotopes facultative in vegetative filaments, and absent in hormogonia. Thylakoids arranged peripherally and parallel to the cell wall. Motility facultative (Komárek et al., 2014; Komárek & Anagnostidis, 2005).

### Cytotoxicity of Th. longitrichum sp. nov. fractions

The human HaCat keratinocytes were used to predict the biocompatibility of *Th. longitrichum* sp. nov. fractions. Cells viability was accessed after 24h and 48h of exposition to increasing concentrations (12.5-200μg mL⁻¹) of aqueous and acetonic fractions, through the MTT viability assay.

HaCat cells viability was not affected after 24 h of exposure to the acetonic fraction, forconcentrations up to 100 μg mL⁻¹ (*p* > 0.05), whereas 200 μg mL⁻¹ induced a significant decrease in cell viability (*p* < 0.05) After 48 h, a clearer concentration-dependent pattern was observed: 100 and 200 μg mL⁻¹ significantly reduced viability (*p* < 0.05), while lower concentrations (≤50 μg mL⁻¹) remained statistically comparable to the control (Fig. S3A). These results indicate that cytotoxicity of the acetonic fraction is both dose- and time-dependent, becoming evident only at higher concentrations and prolonged exposure. In accordance with ISO 10993:5-2009, demonstrating non-cytotoxicity is indicated by achieving a cell viability of at least 70% following exposure (ISO 10993-5, 2009). Thus, the present results indicate the absence of cytotoxicity, despite the significant differences observed in the two highest concentrations.

In contrast, the aqueous fraction displayed a more pronounced early effect. After 24 h of exposure, all tested concentrations (12.5–200 μg mL⁻¹) significantly reduced viability relative to the control (*p* < 0.01), although the magnitude of reduction was moderate and did not follow a clear dose–response trend. After 48 h, lower concentrations (12.5–50 μg mL⁻¹) no longer differed significantly from the control (*p* > 0.05), whereas 100 and 200 μg mL⁻¹ maintained significant reductions in viability (*p* < 0.05) (Fig. S3B), suggesting partial recovery or cellular adaptation at submaximal doses over time.

Additionally, PCR screening was conducted to detect genes involved in the biosynthesis of major cyanotoxins. The presence of mcyE, sxtG, anaC, and cyrA, associated with the production of microcystin, saxitoxin, anatoxin-a, and cylindrospermopsin, respectively, was tested. However, none of these genes were amplified in ***Th. longitrichum* sp. nov. LEGE 10371**, suggesting that this strain lacks the genetic potential to produce these cyanotoxins (Fig. S4).

### Chemical characterization

#### Carotenoid and Chlorophylls Profiling by HPLC-PDA

HPLC-PDA analysis of the acetone fraction of *Th. longitrichum* sp. nov. LEGE 10371 revealed the presence of eleven pigments, including eight carotenoids, chlorophyll-a, and two chlorophyll-a derivatives, as shown in Figure 8. The identified carotenoids comprised three xanthophylls: lutein (1), zeaxanthin (4), and mixoxanthophyll (6), and one carotene, β-carotene (10). Chlorophyll-a (3) was also detected. In addition, four carotenoid-related compounds were observed but could not be identified based solely on UV–Vis spectra and retention time comparison: three additional β-carotene derivatives (7,8,9,11), and two chlorophyll-a derivatives (2, 5). These unidentified compounds displayed absorption spectra consistent with carotenoid and chlorophyll standards, however their retention times did not match those of the available reference compounds, preventing definitive identification.

**Figure 8.**
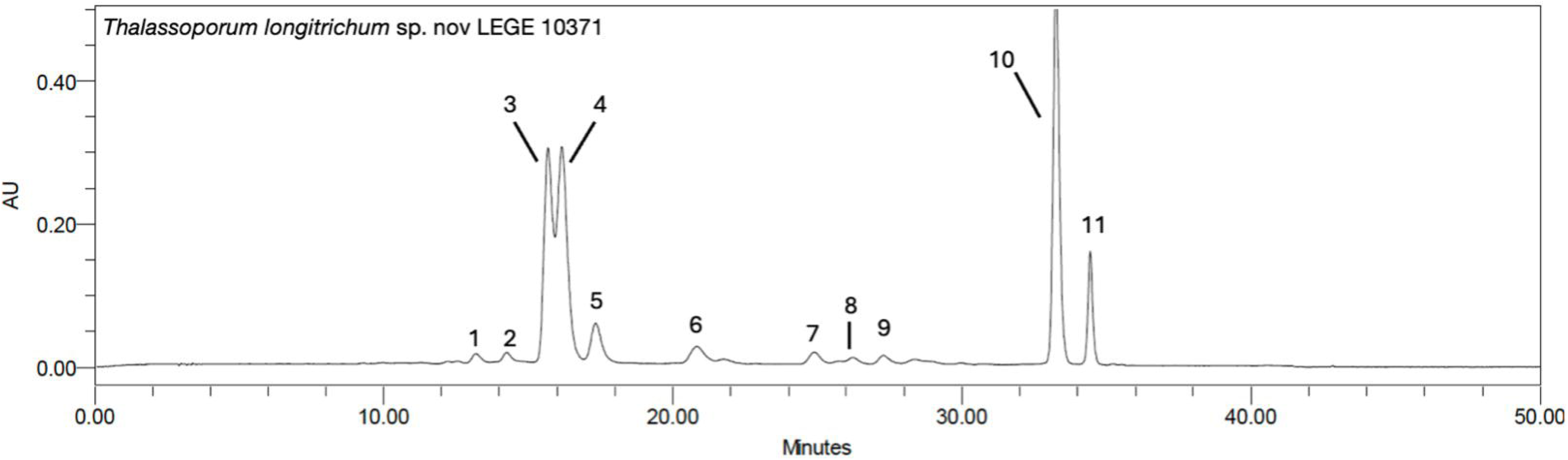
Carotenoid and chlorophylls profile of the acetonic fraction of *Thalassoporum longitrichum* sp. nov. LEGE 10371. HPLC-PDA recorded at 450 nm. Lutein (1), chlorophyll a derivatives (2,5), zeaxanthin (4), chlorophyll a (3), mixoxanthophyll (6), β-carotene derivatives (7,8,9,11), β-carotene (10).

Chlorophyll-a (3) was the most abundant pigment, reaching 216.90 ± 0.13 µg mg⁻¹ dry fraction, equivalent to 2.96 ± <0.01 mg g⁻¹ of cyanobacterial biomass. Among carotenoids, mixoxanthophyll reached 49.55 ± 0.67 µg mg⁻¹ dry fraction (0.68 ± 0.01 mg g⁻¹), while zeaxanthin was quantified at 18.52 ± 0.14 µg mg⁻¹ dry fraction (0.25 ± <0.01 mg g⁻¹). β-Carotene was present at 43.52 ± 0.23 µg mg⁻¹ dry fraction (0.59 ± <0.01 mg g⁻¹) and lutein was present at 1.37 ± 0.06 µg mg⁻¹ dry fraction (0.02 ± <0.01 mg g⁻¹). The quantification of the other pigments are displayed in Table 3. In quantitative terms, chlorophylls accounted for the largest proportion of the pigments, followed by β-carotene, and mixoxanthophyll.

**Table 3.** Carotenoid and chlorophylls content in the acetone fraction of *Thalassoporum longitrichum* sp. nov. LEGE 10371, determined by HPLC-PDA.

| Peak | RT | Compound | Concentration<br>($\mu\text{g}/\text{mg}_{\text{dry fraction}} \pm \text{SD}$) | Concentration<br>($\mu\text{g}/\text{mg}_{\text{dry biomass}} \pm \text{SD}$) |
| --- | --- | --- | --- | --- |
| 1 | 13.12 | Lutein | $1.37 \pm 0.06$ | $0.02 \pm \leq 0.01$ |
| 2 | 14.26 | Chlorophyll a derivative | $29.28 \pm 0.50$ | $0.40 \pm 0.01$ |
| 3 | 15.68 | Chlorophyll-a | $216.90 \pm 0.13$ | $2.96 \pm \leq 0.01$ |
| 4 | 16.16 | Zeaxanthin | $18.52 \pm 0.14$ | $0.25 \pm \leq 0.01$ |
| 5 | 17.33 | Chlorophyll-a derivative | $71.32 \pm 2.45$ | $0.97 \pm 0.03$ |
| 6 | 20.88 | Mixoxanthophyll | $49.55 \pm 0.67$ | $0.68 \pm 0.01$ |
| 7 | 24.88 | β-Carotene derivative | $2.94 \pm 1.41$ | $0.04 \pm 0.02$ |
| 8 | 26.21 | β-Carotene derivative | $1.53 \pm 0.05$ | $0.02 \pm \leq 0.01$ |
| 9 | 27.27 | β-Carotene derivative | $1.61 \pm 0.20$ | $0.02 \pm \leq 0.01$ |
| 10 | 33.26 | β-Carotene | $43.52 \pm 0.23$ | $0.59 \pm \leq 0.01$ |
| 11 | 34.43 | β-Carotene derivative | $4.86 \pm 0.02$ | $0.07 \pm \leq 0.01$ |

#### Phycobiliproteins (PBPs)

Phycobiliproteins, being water-soluble proteins, were quantified exclusively in the aqueous extract. Three main PBPs were detected: phycocyanin (PC), allophycocyanin (APC), and phycoerythrin (PE) (Fig 9). Quantification was performed using modified spectrophotometric equations that minimize chlorophyll-a interference, thereby minimizing spectral overlap and ensuring more accurate estimation of PBP concentrations (Lauceri et al., 2018).

**Figure 9.**
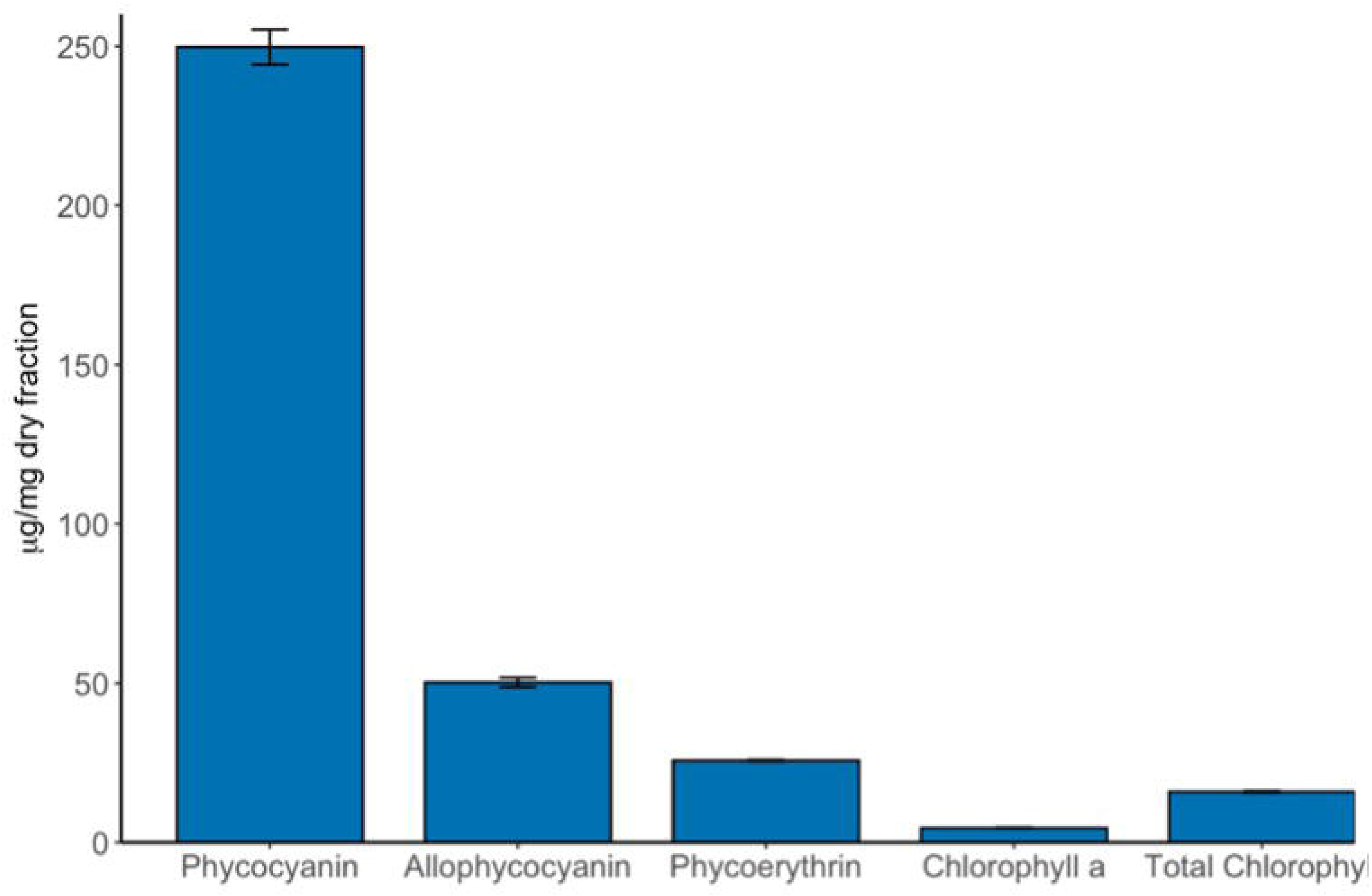
Phycobiliproteins (allophycocyanin, phycocyanin and phycoerythrin) and chlorophylls quantified in the aqueous fraction of *Thalassoporum longitrichum* sp. nov. LEGE 10371.

PC was the predominant phycobiliprotein, reaching 249.73 µg mg⁻¹ dry fraction. APC was quantified at 50.27 µg mg⁻¹ dry fraction, whereas PE was detected at 25.73 µg mg⁻¹ dry fraction. As seen in Figure 9, PC represented the largest fraction of the total PBPs, with APC and PE contributing in smaller proportions. Chlorophyll-a was quantified at 4.50 µg mg⁻¹ dry extract, and total chlorophylls content reached 15.92 µg mg⁻¹ dry extract.

Overall, PBPs were present at substantially higher concentrations than chlorophyll-a in the aqueous fraction. Variability among replicates was low, as indicated by the small error bars observed.

### Bioactivity Assays

#### O_2_^•−^ Scavenging

The aqueous fraction was further evaluated for O_₂_^•⁻^ scavenging, as a complementary indicator of oxidative stress associated with inflammatory processes (Fig. 10A). A concentration-dependent response was observed between 0.013 and 1.667 mg dry fraction mL⁻¹. At 0.013 mg mL⁻¹, radical scavenging was approximately 12%, increasing to ∼30% at 0.052 mg mL⁻¹. A plateau between 30–33% inhibition was detected at intermediate concentrations (0.104–0.833 mg mL⁻¹). The highest tested concentration (1.667 mg mL⁻¹) resulted in approximately 40% scavenging activity. Interpolation analysis using linear regression of the dose–response curve yielded an IC₂₅ value of 0.042–0.045 mg mL⁻¹. In contrast, acetonic fractions did not exhibit measurable O_₂_^•⁻^ scavenging activity within the same concentration range.

**Figure 10.**
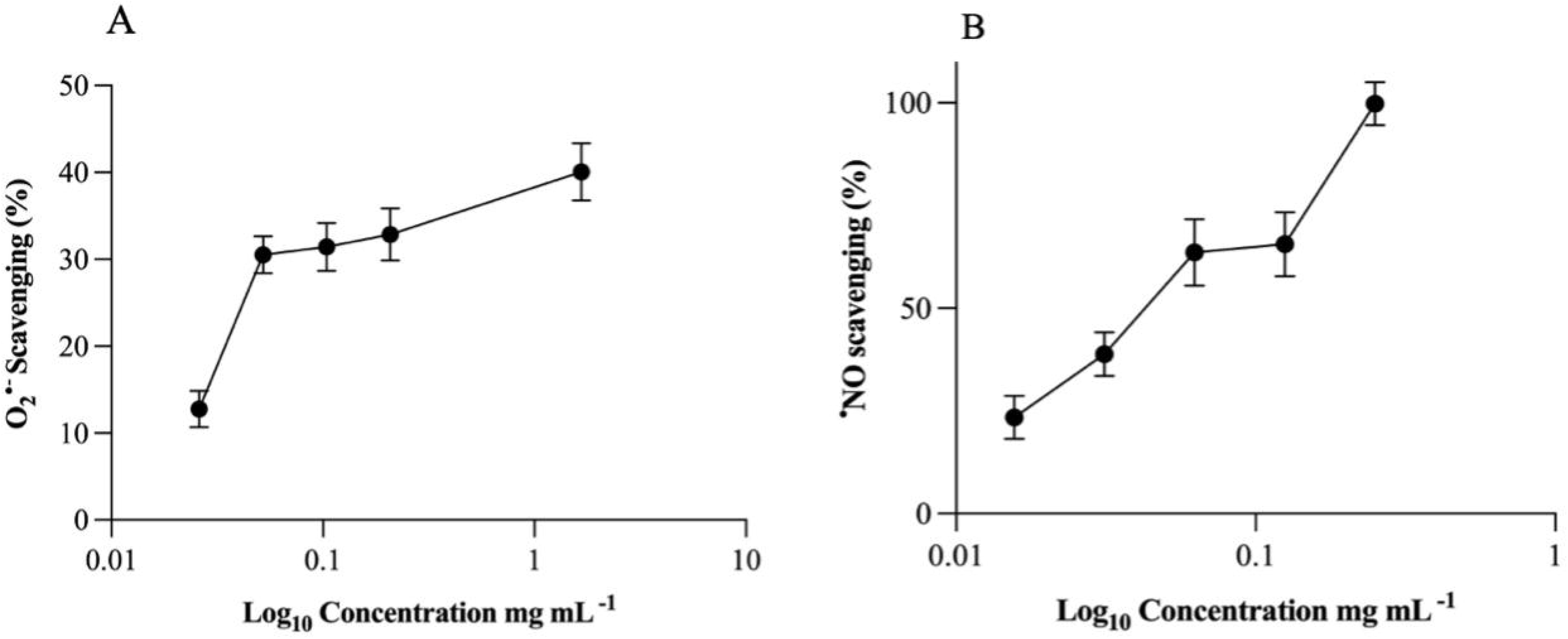
Superoxide anion radical (O_2_^•−^) (A) and nitric oxide radical (^•^NO) (B) scavenging activity of aqueous and acetonic fractions of *Thalassoporum longitrichum* sp. nov. LEGE 10371, respectively. Values are expressed as Log_10_ of the mean ± SE in mg mL^-1^ of dry fraction of at least three independent experiments, performed in duplicate.

#### • NO Scavenging

The anti-inflammatory potential of *Th. longitrichum* sp. nov. LEGE 10371 fractions was predicted by measuring their ability to scavenge ^•^NO, an important mediator involved in vasodilation and recruitment of inflammatory cytokines. A clear concentration-dependent • NO scavenging was observed across the tested range (approximately 0.02–0.25 mg mL⁻¹) (Fig. 10B). At the lowest tested concentration (∼0.02 mg mL⁻¹), ^•^NO scavenging was approximately 20–25%. Scavenging activity increased to ∼40% at 0.04 mg mL⁻¹ and exceeded 60% at 0.08 mg mL⁻¹. At 0.12 mg mL⁻¹, scavenging reached approximately 65–70%, while the highest tested concentration (0.25 mg mL⁻¹) resulted in nearly complete • NO scavenging (∼100%). The fraction reached a value of IC₅₀=0.045 mg mL⁻¹. For comparative purposes, quercetin, used as a reference compound under the same experimental conditions, exhibited an IC₅₀=0.58 mg mL⁻¹. The substantially lower IC₅₀ obtained for the fraction indicates a markedly higher ^•^NO scavenging efficiency in this *in vitro* system.

#### LOX inhibition

The acetonic fraction from *Th. longitrichum* sp. nov. LEGE 10371 showed a moderate inhibitory effect on the enzyme. Under the tested conditions, the extract produced a maximum inhibition of LOX activity of 41.5 ± 12.1, for a fraction concentration of 250 µg/mL, when compared with the untreated control (Figure 11).

**Figure 11.**
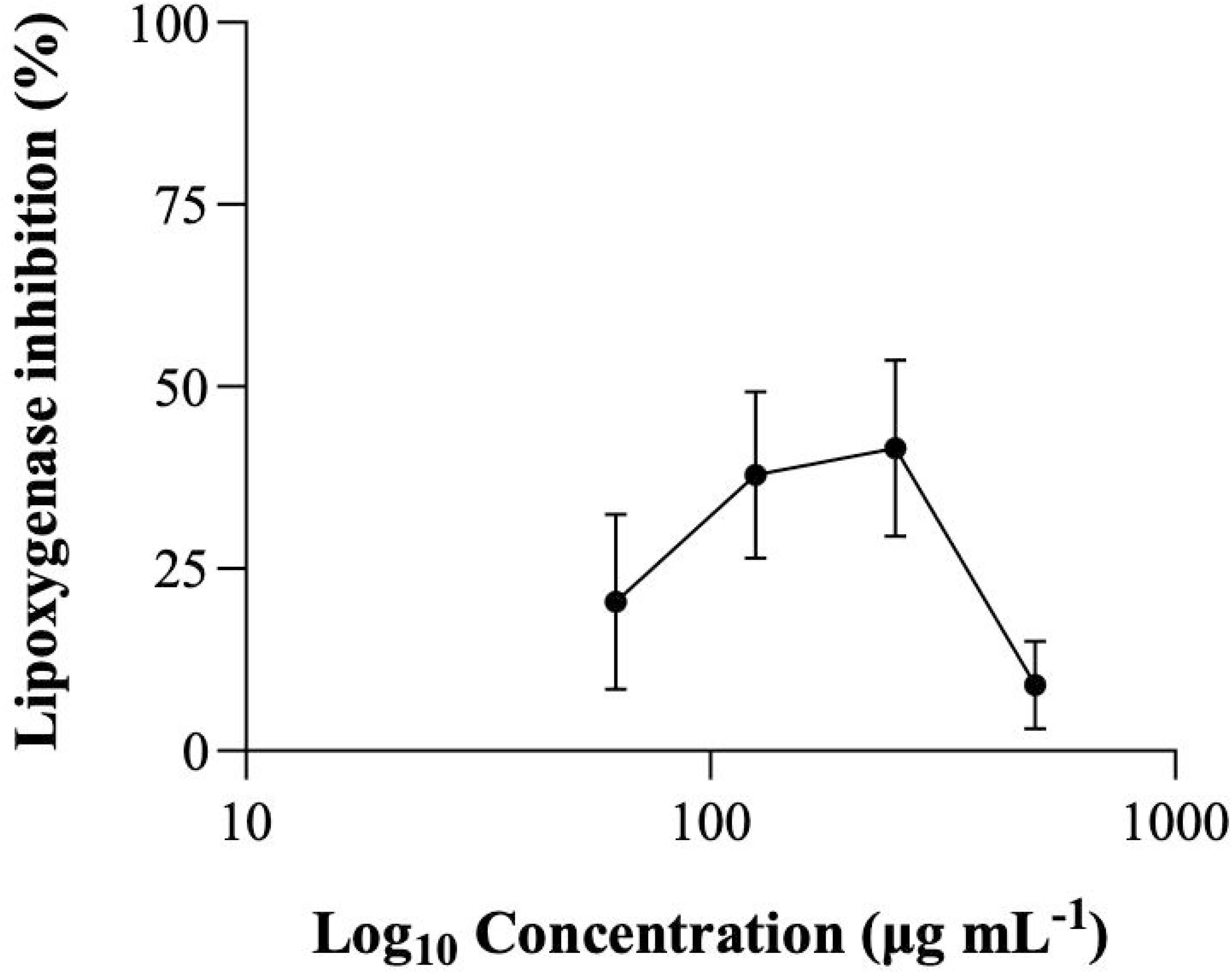
Lipoxygenase inhibition (%) of the acetonic fraction of *Thalassoporum longitrichum* sp. nov. LEGE 10371. Values are expressed as Log_10_ of the mean ± SE in µg mL^-1^ of dry fraction, of at least three independent experiments, performed in duplicate.

## DISCUSSION

The separation of *Pseudanabaena* and *Limnothrix* has long been sustained by subtle morphological distinctions, primarily trichome constriction and the presence or absence of aerotopes (Komárek & Anagnostidis, 2005). However, even within that morphological framework, these generic boundaries were acknowledged as unstable. Species such as *P. galeata* SAG 13.83 and *P. pruinosa* have been reported to possess aerotopes, traditionally considered diagnostic for *Limnothrix*, illustrating the morphological overlap between both genera (Aleksovski et al., 2024).

The molecular data presented here confirms that instability. Strains attributed to *Limnothrix*, including representatives of the type species *L. redekei*, cluster unequivocally within the *Pseudanabaena* lineage, forming a single, well-supported clade. Similar patterns have been reported in previous phylogenetic studies, where *L. redekei* strains show extremely high sequence similarity with *Pseudanabaena* isolates (Akagha et al., 2019; Luz et al., 2023; Strunecký et al., 2023; Saraf et al., 2025).

The exceptionally high 16S rRNA gene identity values observed between strains assigned to both genera, reaching up to 99.9% are incompatible with generic separation under modern standards. Yarza et al. (2014) proposed a 94.5% sequence identity threshold for genus-level delineation. The values recovered here fall well above that boundary. Comparable phylogenetic intermixing has been reported in freshwater ecosystems, where morphologically defined *Pseudanabaena* and *Limnothrix* populations occupy also overlapping ecological niches and cluster together in molecular surveys (Acinas et al., 2008; Akagha et al., 2019; Gkelis et al., 2005). Additionally, the D1-D1’ helices of the 16S-23S ITS region were very similar among *Pseudanabaena/“Limnothrix”* strains with a consistent UBB region.

Ecologically, both lineages share opportunistic strategies typical of filamentous cyanobacteria inhabiting eutrophic environments. They tolerate broad gradients of light and nutrient availability and frequently contribute to dominant populations under nutrient enrichment (Acinas et al., 2008; Gkelis et al., 2005; Paerl & Otten, 2013; Zhu et al., 2012). Both have the ability to thrive in fluctuating, organic-rich waters which likely contributes to their frequent coexistence in lakes, estuaries, and coastal systems (Acinas et al., 2008). No consistent ecological partitioning has been demonstrated between them, further weakening the basis for generic separation.

Based on these evidences, the concept that *Pseudanabaena* and *Limnothrix* represent the same generic entity is not new. However, no formal taxonomic proposal has been made so far to unite these genera. Our study validates these previous observations and, accordingly, formally proposes the synonymization of *Limnothrix* under *Pseudanabaena*.

In contrast, *Thalassoporum* forms a distinct and well-supported lineage. Strain LEGE 10371 clusters within this genus but is clearly separated from *Th. komarekii* and *Th. mexicanum*. The <98% 16S rRNA identity values with the other species of the genus support its recognition as a novel species. According to Yarza et al. (2014), 16S rRNA gene similarity values below 98.7% are strong indicative to strains belong to different species. Additionally, the marked divergence in D1–D1′ helix configuration confirm the separation of the three species of the genus. Morphological traits, including longer trichomes and the absence of aerotopes, further distinguish *Th. longitrichum* sp. nov. from its congeners.

While the primary focus of Pseudanabaenales research has been taxonomic resolution (Komárek et al., 2014; (Strunecký et al., 2023) increasing attention has been directed toward the pigment composition and associated bioactivities of filamentous cyanobacteria (Favas et al., 2022; Pagels et al., 2021). In the case of the pigment composition of *Th. longitrichum* sp. nov. reveals a metabolically diverse profile dominated by chlorophylls, carotenoids (β-carotene, zeaxanthin, lutein, mixoxanthophyll), and substantial phycobiliprotein content. In cyanobacteria, these pigments play fundamental roles in photosynthesis and photoprotection but have also been linked to antioxidant and anti-inflammatory properties (Duarte et al., 2026; Favas et al., 2022).

The functional assays conducted herein indicate that the chemical diversity of cyanobacteria fractions is associated with measurable radical scavenging activity in cell-free systems. The aqueous fraction, enriched in PBPs, exhibited concentration-dependent ability to scavenge O_₂_^•⁻^. Superoxide represents a primary reactive oxygen species involved in inflammatory oxidative cascades, and its modulation is frequently used as an indicator of antioxidant capacity in natural extracts. PBPs, particularly PC, has been reported to scavenge reactive oxygen species and to attenuate oxidative damage *in vitro* (Benedetti et al., 2004; Pagels et al., 2021; Romay et al., 1998).

In contrast, the acetonic fraction demonstrated significant ^•^NO scavenging activity, reaching a value of IC₅₀=45μg mL^-1^. ^•^NO is a key mediator of inflammatory signaling; while physiologically essential, its uncontrolled production contributes to nitrosative stress and tissue damage. Carotenoids are known to interact with reactive nitrogen species and to modulate redox-sensitive pathways(Pagels et al., 2021). Moreover, pigment-rich cyanobacterial extracts have also been shown to influence pro-inflammatory enzymes such as LOX and cyclooxygenase (COX), leading to the reduction of inflammatory mediators (Rodrigues et al., 2024). In the present study, the acetonic fraction of *Th. longitrichum* sp. nov. LEGE 10371 exhibited a moderate inhibition of LOX activity. This suggests that the bioactive compounds present in this fraction may also have potential to inhibit the production of leukotrienes, which are important molecules involved in the exacerbation of the inflammatory process (Rodrigues et al., 2024).

The differential chemical profile between aqueous and acetonic fractions, derived from the polarity-dependent extraction of distinct pigment classes, is reflected in their targeted ability to scavenge specific free radicals. This is an advantage of the sequential extraction procedure, once it allows the creation of a more precise, function-specific antioxidant system, through the production of fractions with distinct targets from the same biomass. Moreover, this feature allows the combination or strategical dosage of the fractions aiming higher efficiency and expanded biotechnological applications.

It is worth highlight the complementarity of ^•^NO and O_₂_^•⁻^ in the inflammatory process. NO reacts with superoxide to form peroxynitrite (ONOO^−^), another reactive nitrogen species that, together with NO, function not only as toxic but also as intracellular signalling molecules. These redox dynamics can simultaneously regulate transcription factors, such as NF-κB, which act as a key mechanism for determining the pro-inflammatory or anti-inflammatory polarization states of macrophages, by reprogramming inflammatory genes expression and cellular metabolism (Morgan & Liu, 2010; Radi, 2013). However, it is important to mention that anti-inflammatory interpretation of ^•^NO scavenging must be approached cautiously. *In vitro* radical scavenging does not necessarily equate to modulation of cellular inflammatory pathways, and its biological relevance requires validation in cellular models.

Nevertheless, NO inhibition assays are widely used as preliminary indicators of anti-inflammatory potential in cyanobacterial and microalgal extracts, particularly when combined with pigment characterization (Favas et al., 2022; Morone et al., 2022; Romay et al., 1998). The fact that ^•^NO scavenging was detectable at relatively low concentrations suggests that the carotenoid-rich fraction of *Th. longitrichum* sp. nov. LEGE 10371 may contribute to redox modulation under inflammatory conditions, although mechanistic studies would be required to confirm interactions with enzymes such as inducible nitric oxide synthase (iNOS).

Equally relevant is the cytotoxicity assessment. It constitutes an essential component in the characterization of natural extracts, allowing to predict potential harmful effects that could limit their suitability for health-related applications. Moreover, the use of *in vitro* cytotoxicity models aligns with current efforts to reduce animal experimentation, promotes a more sustainable and ethically responsible research framework, and strengthens the evidence supporting the potential incorporation of these fractions into pharmaceutical, cosmetic, or nutraceutical formulations. In this context, both the acetonic and aqueous fractions appear to demonstrate a safe profile within the concentration range of 12.5–50 μg mL⁻¹, where no marked or persistent reduction in cells viability was observed. Nevertheless, other cell lines are worth of further exploitation in order to get a broader picture of the security profile.

Contemporary cyanobacterial systematics increasingly integrates molecular phylogeny with phenotypic and biochemical traits (Duarte et al., 2026). Comparative genomic studies have demonstrated that lineage diversification within Cyanobacteria often coincides with substantial metabolic differentiation, particularly in secondary metabolite biosynthetic capacity (Calteau et al., 2014; Shih et al., 2013). The integration of phylogenetic resolution, ITS structural analysis, morphology, ecology, and metabolites profiling, may thus illustrate how evolutionary coherence can be evaluated alongside functional attributes within a modern polyphasic framework. Unfortunately, due to the total absence of studies focused exclusively on the biochemical profiling of *Thalassoporum* sp. pigments, it is not yet possible to establish a robust correlation.

Taken together, the synonymization of *Limnothrix* under *Pseudanabaena* is supported by convergent molecular and ecological evidence, whereas *Thalassoporum* remains a distinct marine lineage (Strunecký et al., 2023). The characterization of *Th. longitrichum* sp. nov. expands both the taxonomic and functional landscape of Thalassoporaceae, demonstrating that systematic clarification and metabolic profiling can be cohesively addressed within integrative cyanobacterial research.

## Supporting information

Supplemental Figure 1. 16S rRNA gene phylogenetic tree with 436 sequences.

Supplemental Figure 2. MALDI-TOF MS spectrum

Supplemental Figure 3. Cytotoxic effects of acetonic extracts (A) and aqueous (B) fractions in HaCaT keratinocytes at 24 and 48h.

Supplemental Figure 4. PCR screening for cyanotoxin

Supplemental Table 1. Table S1. 16S rRNA gene identity between Thalassoporum strains.

Supplemental Table 2. 16S rRNA gene sequence identity between Limnothrix and Pseudanabaena

## Acknowledgements

Flavio Oliveira thanks to National funds through FCT scholarship grant UI/BD/06241/2021. Graciliana Lopes thanks FCT for the financial support through the Scientific Employment Stimulus-Individual Call (https://doi.org/10.54499/2021.01768.CEECIND/CP1665/CT0007). The MALDI-TOF MS profiles were obtained by Célia Maria Gonçalves Soares and Sónia Raquel Alves Fernandes Pereira, University of Minho.

## Supplementary Figures

Figure S1. 16S rRNA gene phylogenetic tree with 436 sequences. Bootstrap values above 90% are presented at the nodes. The Pseudanabaenales are marked with the dark green strip and *Thalassoporum* clade is shaded in light green.

Figure S2. MALDI-TOF MS spectrum of strain *Th. longitrichum* sp. nov. LEGE 10731 acquired in the m/z range of 2,000–20,000, showing the main protein.

Figure S3. Cytotoxic effects of acetonic extracts (A) and aqueous (B) fractions in HaCaT keratinocytes at 24 and 48h.

Figure S4. PCR screening for cyanotoxin biosynthesis genes in *Th. longitrichum* sp. nov. LEGE 10371. (A) *cyrA* gene associated with cylindrospermopsin biosynthesis; (B) *sxtG* gene associated with saxitoxin biosynthesis; (C) *mcyE* gene associated with microcystin biosynthesis; (D) *anaC* gene associated with anatoxin biosynthesis. Lanes correspond to NC (negative control), PC (positive control), strain LEGE 10371, and the DNA ladder.

Table S1. 16S rRNA gene identity between *Thalassoporum* strains.

Table S2. 16S rRNA gene sequence identity between *Limnothrix* and *Pseudanabaena*

## References

Acinas, S. G., Haverkamp, T. H. A., Huisman, J., & Stal, L. J. (2008). Phenotypic and genetic diversification of Pseudanabaena spp. (cyanobacteria). The ISME Journal 2009 3:1, 3(1), 31–46. 10.1038/ismej.2008.78

Akagha, S. C., Johansen, J. R., Nwankwo, D. I., & Yin, K. (2019). Lagosinema tenuis gen. et sp. nov. (Prochlorotrichaceae, Cyanobacteria): a new brackish water genus from Tropical Africa. Fottea, 19(1), 1–12. 10.5507/fot.2018.012

Aleksovski, B., Krstić, S., Komárek, J., Nguyen, K., Pakovski, K., Kiprijanovska, S., Dimovski, A., Vuchurević, A., Stefanoska, E., & Strunecký, O. (2024). Pseudanabaena pruinosa sp. nov. (Pseudanabaenales, Cyanobacteria): an Arctic Pseudanabaena species with branched sheaths and central aerotopes. European Journal of Phycology, 59(3), 311–331. 10.1080/09670262.2024.2343088

Anagnostidis, K., & Komárek, J. (1988). Modern approach to the classification system of cyanophytes. 3 – Oscillatoriales. Algological Studies, 50–53, 327–472.

Benedetti, S., Benvenuti, F., Pagliarani, S., Francogli, S., Scoglio, S., & Canestrari, F. (2004). Antioxidant properties of a novel phycocyanin extract from the blue-green alga Aphanizomenon flos-aquae. Life Sciences, 75(19), 2353–2362. 10.1016/j.lfs.2004.06.004

Bennett, A., & Bogobad, L. (1973). Complementary chromatic adaptation in a filamentous blue-green alga. The Journal of Cell Biology, 58(2), 419–435. 10.1083/jcb.58.2.419

Benredjem, L., Morais, J., Hentschke, G. S., Abdi, A., Berredjem, H., & Vasconcelos, V. (2023). First Polyphasic Study of Cheffia Reservoir (Algeria) Cyanobacteria Isolates Reveals Toxic Picocyanobacteria Genotype. Microorganisms 2023, Vol. 11, 11(11). 10.3390/microorganisms11112664

Bornet, J.-B. É., & Flahault, C. (1886). Révision des Nostocacées hétérocystées contenues dans les principaux herbiers de France. Annales des Sciences Naturelles, Botanique, 7, 3, 323–381.

Bornet, J.-B. É., & Flahault, C. (1888). Révision des Nostocacées hétérocystées contenues dans les principaux herbiers de France. Annales des Sciences Naturelles, Botanique, 7, 7, 177–262.

Calteau, A., Fewer, D. P., Latifi, A., Coursin, T., Laurent, T., Jokela, J., Kerfeld, C. A., Sivonen, K., Piel, J., & Gugger, M. (2014). Phylum-wide comparative genomics unravel the diversity of secondary metabolism in Cyanobacteria. BMC Genomics 2014 15:1, 15(1), 977-. 10.1186/1471-2164-15-977

Duarte, A. J. C., Hentschke, G. S., Oliveira, F., Vasconcelos, V., & Lopes, G. (2026). Description of a New Marine Cyanobacterium from the Cabo Verde Archipelago: Pigments Profile and Biotechnological Potential of Salileptolyngbya caboverdiana sp. nov. Marine Drugs 2026, Vol. 24, 24(1), 29. 10.3390/md24010029

Favas, R., Morone, J., Martins, R., Vasconcelos, V., & Lopes, G. (2022). Cyanobacteria Secondary Metabolites as Biotechnological Ingredients in Natural Anti-Aging Cosmetics: Potential to Overcome Hyperpigmentation, Loss of Skin Density and UV Radiation-Deleterious Effects. Marine Drugs 2022, Vol. 20, 20(3). 10.3390/md20030183

Fernandes, F., Barbosa, M., Pereira, D. M., Sousa-Pinto, I., Valentão, P., Azevedo, I. C., & Andrade, P. B. (2018). Chemical profiling of edible seaweed (Ochrophyta) extracts and assessment of their in vitro effects on cell-free enzyme systems and on the viability of glutamate-injured SH-SY5Y cells. Food and Chemical Toxicology, 116(Pt B), 196–206. 10.1016/j.fct.2018.04.033

Gkelis, S., Rajaniemi, P., Vardaka, E., Moustaka-Gouni, M., Lanaras, T., & Sivonen, K. (2005). Limnothrix redekei (Van Goor) Meffert (Cyanobacteria) Strains from Lake Kastoria, Greece Form a Separate Phylogenetic Group. Microbial Ecology 2005 49:1, 49(1), 176–182. 10.1007/s00248-003-2030-7

Gomont, M. (1892). Monographie des Oscillariées (Nostocacées Homocystées). Deuxième partie. - Lyngbyées. Annales Des Sciences Naturelles, Botanique, 7(16), 91–264.

Hentschke, G., de Oliveira, F., Morais, J., & Vasconcelos, V. (2025). New terrestrial cyanobacterial species from the vicinity of the CIIMAR building on the Portuguese Coast. European Journal of Phycology. 10.1080/09670262.2025.2564078;ISSUE:ISSUE:DOI

Jungblut, A. D., & Neilan, B. A. (2006). Molecular identification and evolution of the cyclic peptide hepatotoxins, microcystin and nodularin, synthetase genes in three orders of cyanobacteria. Archives of Microbiology 2006 185:2, 185(2), 107–114. 10.1007/s00203-005-0073-5

Katoh, K., Misawa, K., Kuma, K. I., & Miyata, T. (2002). MAFFT: a novel method for rapid multiple sequence alignment based on fast Fourier transform. Nucleic Acids Research, 30(14), 3059–3066. 10.1093/nar/gkf436

Komárek, J., & Anagnostidis, K. (1986). Modern approach to the classification system of cyanophytes. 2 – Chroococcales. Algological Studies, 43, 157–226.

Komárek, J., & Anagnostidis, K. (2005). Cyanoprokaryota 2.Teil: Oscillatoriales. Elsevier GmbH.

Komárek, J., Kaštovský, J., Mareš, J., & Johansen, J. R. (2014). Taxonomic classification of cyanoprokaryotes (cyanobacterial genera) 2014, using a polyphasic approach. Preslia, 86, 295–335.

Konstantinou, D., Voultsiadou, E., Panteris, E., & Gkelis, S. (2021). Revealing new sponge-associated cyanobacterial diversity: Novel genera and species. Molecular Phylogenetics and Evolution, 155. 10.1016/j.ympev.2020.106991

Kotai, J. (1972). Instructions for Preparation of Modified Nutrient Solution Z8 for Algae. Norwegian Institute for Water Research.

Lane, D. J. (1991). 16S/23S rRNA Sequencing. In E. Stackebrandt & M. Goodfellow (Eds.), Nucleic Acid Techniques in Bacterial Systematics (pp. 115–175). John Wiley and Sons.

Lauceri, R., Chini Zittelli, G., Maserti, B., & Torzillo, G. (2018). Purification of phycocyanin from Arthrospira platensis by hydrophobic interaction membrane chromatography. Algal Research, 35, 333–340. 10.1016/j.algal.2018.09.003

Leão, P. N., Engene, N., Antunes, A., Gerwick, W. H., & Vasconcelos, V. (2012). The chemical ecology of cyanobacteria. Natural Product Reports, 29(3), 372. 10.1039/c2np00075j

Letunic, I., & Bork, P. (2021). Interactive Tree Of Life (iTOL) v5: an online tool for phylogenetic tree display and annotation. Nucleic Acids Research, 49(W1), W293–W296. 10.1093/nar/gkab301

Lopes, G., Sousa, C., Silva, L. R., Pinto, E., Andrade, P. B., Bernardo, J., Mouga, T., & Valentão, P. (2012). Can phlorotannins purified extracts constitute a novel pharmacological alternative for microbial infections with associated inflammatory conditions? PloS One, 7(2). 10.1371/journal.pone.0031145

Lopes, V. R., Ramos, V., Martins, A., Sousa, M., Welker, M., Antunes, A., & Vasconcelos, V. M. (2012). Phylogenetic, chemical and morphological diversity of cyanobacteria from Portuguese temperate estuaries. Marine Environmental Research, 73, 7–16. 10.1016/j.marenvres.2011.10.005

Lukesova, A., Johansen, J. R., Martin, M. P., & Casamatta, D. A. (2009). Aulosira bohemensis sp. nov.: Further phylogenetic uncertainty at the base of the Nostocales (Cyanobacteria). Phycologia, 48(2), 118–129. 10.2216/08-56.1

Luz, R., Cordeiro, R., Kaštovský, J., Johansen, J. R., Dias, E., Fonseca, A., Urbatzka, R., Vasconcelos, V., & Gonçalves, V. (2023). Description of four new filamentous cyanobacterial taxa from freshwater habitats in the Azores Archipelago. Journal of Phycology, 59(6), 1323–1338. 10.1111/JPY.13396;JOURNAL:JOURNAL:15298817;WGROUP:S TRING:PUBLICATION

Mihali, T. K., Kellmann, R., Muenchhoff, J., Barrow, K. D., & Neilan, B. A. (2008). Characterization of the gene cluster responsible for cylindrospermopsin biosynthesis. Applied and Environmental Microbiology, 74(3), 716–722. 10.1128/AEM.01988-07

Miller, M. A., Pfeiffer, W., & Schwartz, T. (2010). Creating the CIPRES Science Gateway for inference of large phylogenetic trees. Grid Computing Environments. 10.1109/GCE.2010.5676129

Morgan, M. J., & Liu, Z. G. (2010). Crosstalk of reactive oxygen species and NF-κB signaling. Cell Research 2011 21:1, 21(1), 103–115. 10.1038/cr.2010.178

Morone, J., Lopes, G., Morais, J., Neves, J., Vasconcelos, V., & Martins, R. (2022). Cosmetic Application of Cyanobacteria Extracts with a Sustainable Vision to Skincare: Role in the Antioxidant and Antiaging Process. Marine Drugs, 20(12), 761. 10.3390/MD20120761/S1

Napiórkowska-Krzebietke, A., Dunalska, J. A., & Bogacka-Kapusta, E. (2023). Ecological Implications in a Human-Impacted Lake—A Case Study of Cyanobacterial Blooms in a Recreationally Used Water Body. International Journal of Environmental Research and Public Health 2023, Vol. 20, 20(6). 10.3390/ijerph20065063

Neilan, B. A., Jacobs, D., Del Dot, T., Blackall, L. L., Hawkins, P. R., Cox, P. T., & Goodman, A. E. (1997). rRNA sequences and evolutionary relationships among toxic and nontoxic cyanobacteria of the genus Microcystis. International Journal of Systematic Bacteriology, 47(3), 693–697. 10.1099/00207713-47-3-693

693

Nübel, U., Garcia-Pichel, F., & Muyzer, G. (1997). PCR primers to amplify 16S rRNA genes from cyanobacteria. Applied and Environmental Microbiology, 63(8), 3327–3332. 10.1128/aem.63.8.3327-3332.1997

Paerl, H. W., & Otten, T. G. (2013). Harmful Cyanobacterial Blooms: Causes, Consequences, and Controls. Microbial Ecology 2013 65:4, 65(4), 995–1010. 10.1007/S00248-012-0159-Y

Pagels, F., Vasconcelos, V., & Guedes, A. C. (2021). Carotenoids from Cyanobacteria: Biotechnological Potential and Optimization Strategies. Biomolecules, 11(5), 735. 10.3390/BIOM11050735

Price, M. N., Dehal, P. S., & Arkin, A. P. (2009). FastTree: Computing Large Minimum Evolution Trees with Profiles instead of a Distance Matrix. Molecular Biology and Evolution, 26(7), 1641–1650. 10.1093/molbev/msp077

Radi, R. (2013). Peroxynitrite, a stealthy biological oxidant. Journal of Biological Chemistry, 288(37), 26464–26472. 10.1074/jbc.R113.472936

Rantala-Ylinen, A., Känä, S., Wang, H., Rouhiainen, L., Wahlsten, M., Rizzi, E., Berg, K., Gugger, M., & Sivonen, K. (2011). Anatoxin-a Synthetase Gene Cluster of the Cyanobacterium Anabaena sp. Strain 37 and Molecular Methods To Detect Potential Producers. Applied and Environmental Microbiology, 77(20), 7271–7278. 10.1128/AEM.06022-11

Ritchie, R. J. (2008). Universal chlorophyll equations for estimating chlorophylls a, b, c, and d and total chlorophylls in natural assemblages of photosynthetic organisms using acetone, methanol, or ethanol solvents. Photosynthetica 2008 46:1, 46(1), 115–126. 10.1007/s11099-008-0019-7

Rodrigues, L., Morone, J., Scotta Hentschke, G., Vasconcelos, V., & Lopes, G. (2024). Anti-Inflammatory Activity of Cyanobacteria Pigment Extracts: Physiological Free Radical Scavenging and Modulation of iNOS and LOX Activity. 10.3390/md22030131

Rojo, C., & Alvarez Cobelas, M. (1994). Population dynamics of Limnothrix redekei, Oscillatoria lanceaeformis, Planktothrix agardhii and Pseudanabaena limnetica (cyanobacteria) in a shallow hypertrophic lake (Spain). Hydrobiologia, 275–276(1), 165–171. 10.1007/BF00026708

Romay, C., Armesto, J., Remirez, D., González, R., Ledon, N., & García, I. (1998). Antioxidant and anti-inflammatory properties of C-phycocyanin from blue-green algae. Inflammation Research, 47(1), 36–41. 10.1007/s000110050256

Ronquist, F., Teslenko, M., Van Der Mark, P., Ayres, D. L., Darling, A., Höhna, S., Larget, B., Liu, L., Suchard, M. A., & Huelsenbeck, J. P. (2012). MrBayes 3.2: Efficient Bayesian Phylogenetic Inference and Model Choice Across a Large Model Space. Systematic Biology, 61(3), 539–542. 10.1093/sysbio/sys029

Saraf, A., Aleksovski, B., Blondet, E., Krstić, S., Criscuolo, A., & Gugger, M. (2025). Expanding species diversity in the monotypic genera Thalassoporum and Tumidithrix (Pseudanabaenales, Cyanobacteriota) with the description of Thalassoporum mexicanum sp. nov. and Tumidithrix helvetica sp. nov. International Journal of Systematic and Evolutionary Microbiology, 75(8), 006869. 10.1099/ijsem.0.006869

Shih, P. M., Wu, D., Latifi, A., Axen, S. D., Fewer, D. P., Talla, E., Calteau, A., Cai, F., Tandeau De Marsac, N., Rippka, R., Herdman, M., Sivonen, K., Coursin, T., Laurent, T., Goodwin, L., Nolan, M., Davenport, K. W., Han, C. S., Rubin, E. M., … Kerfeld, C. A. (2013). Improving the coverage of the cyanobacterial phylum using diversity-driven genome sequencing. Proceedings of the National Academy of Sciences of the United States of America, 110(3), 1053–1058. 10.1073/pnas.1217107110

Strunecký, O., Ivanova, A. P., & Mareš, J. (2023). An updated classification of cyanobacterial orders and families based on phylogenomic and polyphasic analysis. Journal of Phycology, 59(1), 12–51. 10.1111/jpy.13304

Tamura, K., Stecher, G., & Kumar, S. (2021). MEGA11: Molecular Evolutionary Genetics Analysis Version 11. Molecular Biology and Evolution, 38(7), 3022–3027. 10.1093/molbev/msab120

Taton, A., Grubisic, S., Brambilla, E., De Wit, R., & Wilmotte, A. (2003). Cyanobacterial diversity in natural and artificial microbial mats of Lake Fryxell (McMurdo Dry Valleys, Antarctica): a morphological and molecular approach. Applied and Environmental Microbiology, 69(9), 5157–5169. 10.1128/AEM.69.9.5157-5169.2003

Trifinopoulos, J., Nguyen, L. T., von Haeseler, A., & Minh, B. Q. (2016). W-IQ-TREE: a fast online phylogenetic tool for maximum likelihood analysis. Nucleic Acids Research, 44(W1), W232–W235. 10.1093/NAR/GKW256

Whitton, B. A. (2012). Ecology of Cyanobacteria II: Their Diversity in Space and Time. Springer Science & Business Media.

Yarza, P., Yilmaz, P., Pruesse, E., Glöckner, F. O., Ludwig, W., Schleifer, K. H., Whitman, W. B., Euzéby, J., Amann, R., & Rosselló-Móra, R. (2014). Uniting the classification of cultured and uncultured bacteria and archaea using 16S rRNA gene sequences. Nature Reviews Microbiology 2014 12:9, 12(9), 635–645. 10.1038/nrmicro3330

Zhu, M., Yu, G., Li, X., Tan, W., & Li, R. (2012). Taxonomic and phylogenetic evaluation of Limnothrix strains (Oscillatoriales, Cyanobacteria) by adding Limnothrix planktonica strains isolated from central China. Phytoplankton Responses to Human Impacts at Different Scales, 367–374. 10.1007/978-94-007-5790-5_26

Zuker, M. (2003). Mfold web server for nucleic acid folding and hybridization prediction. Nucleic Acids Research, 31(13), 3406–3415. 10.1093/nar/gkg595

