## Supplementary figures and images for "*Thalassoporum longitrichum* sp. nov., a marine epizoic cyanobacterium with anti-inflammatory potential, and the taxonomic reassessment of *Limnothrix* Meffert"

### Supplemental Figure 1. 16S rRNA gene phylogenetic tree with 436 sequences.

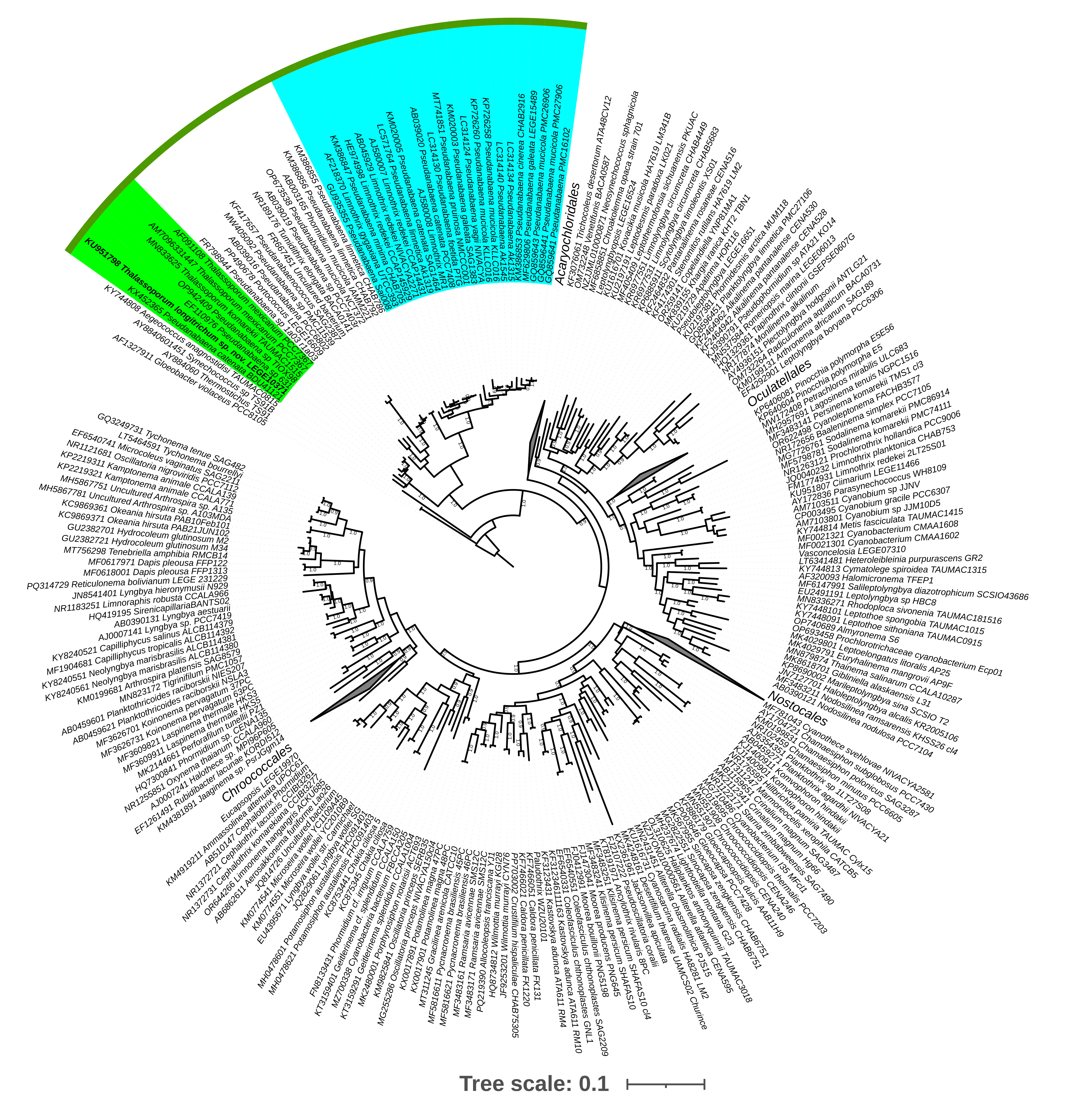

### Supplemental Figure 2. MALDI-TOF MS spectrum

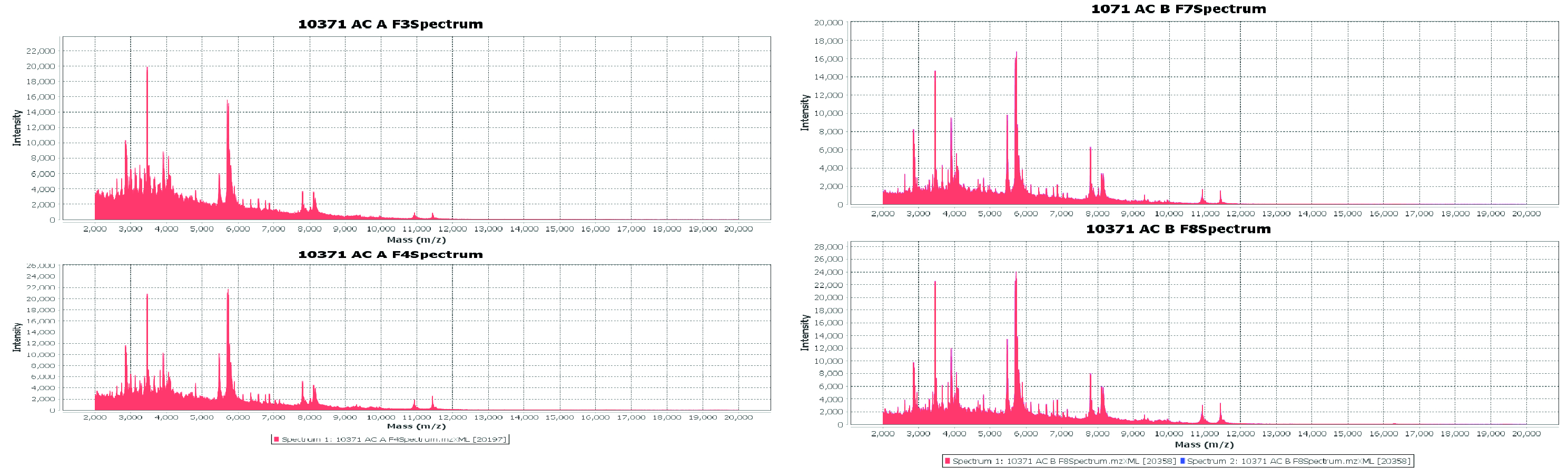

### Supplemental Figure 3. Cytotoxic effects of acetonic extracts (A) and aqueous (B) fractions in HaCaT keratinocytes at 24 and 48h.

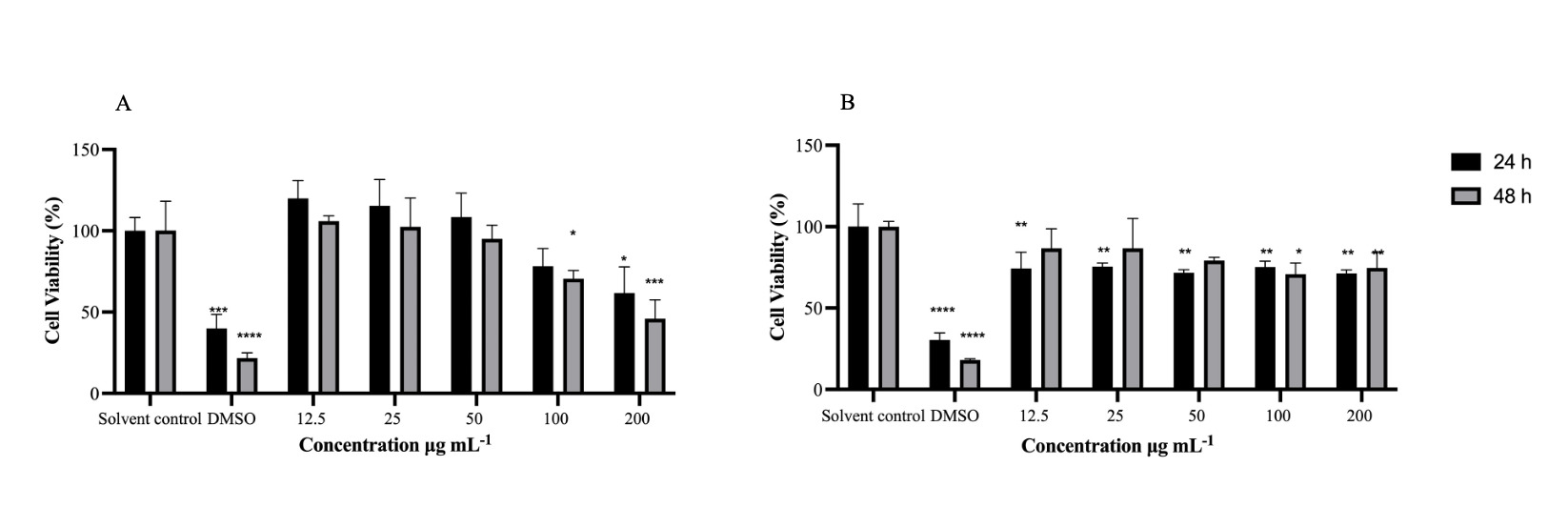

### Supplemental Figure 4. PCR screening for cyanotoxin

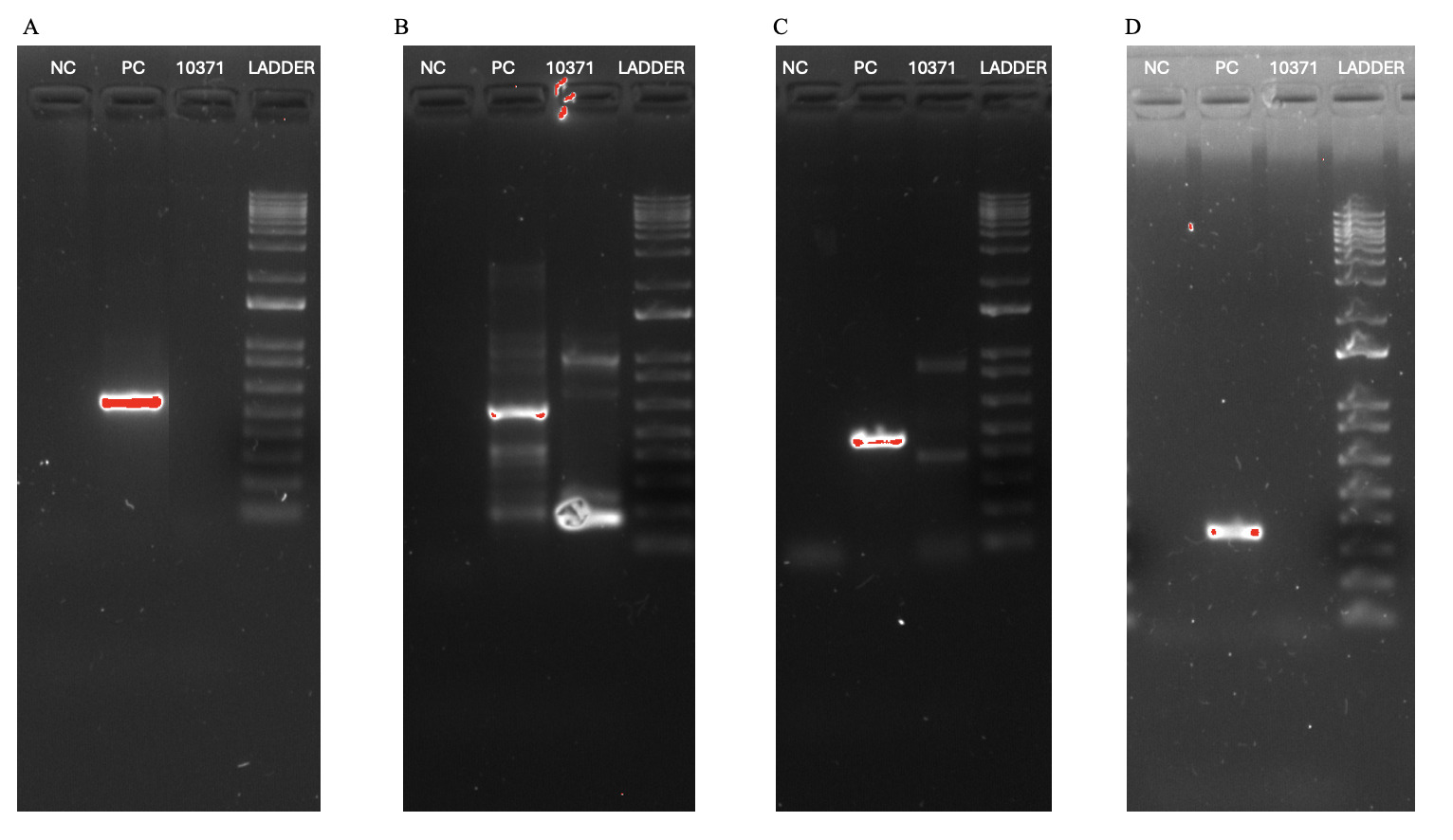

### Supplemental Table 1. Table S1. 16S rRNA gene identity between Thalassoporum strains.

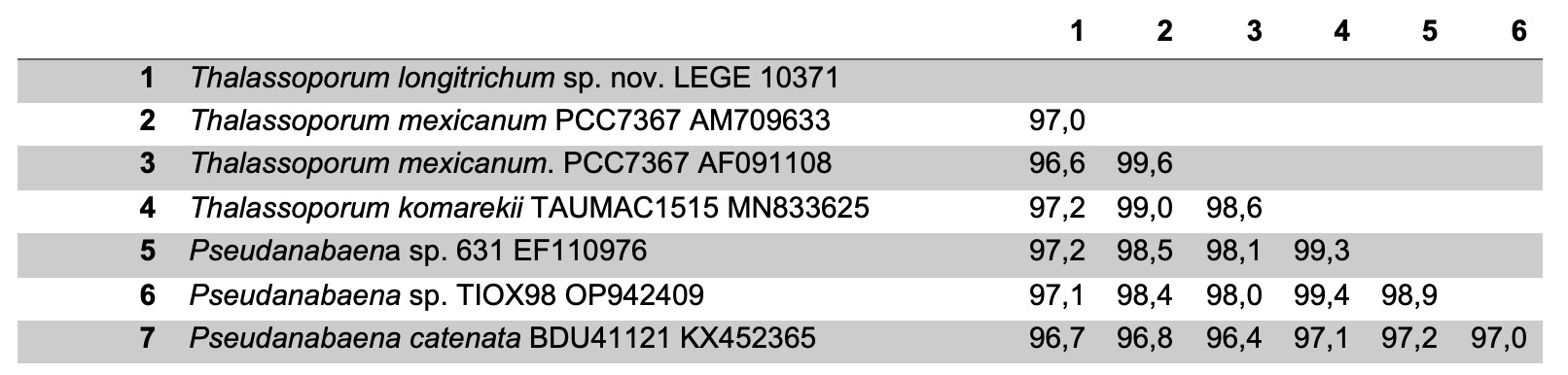

### Supplemental Table 2. 16S rRNA gene sequence identity between Limnothrix and Pseudanabaena

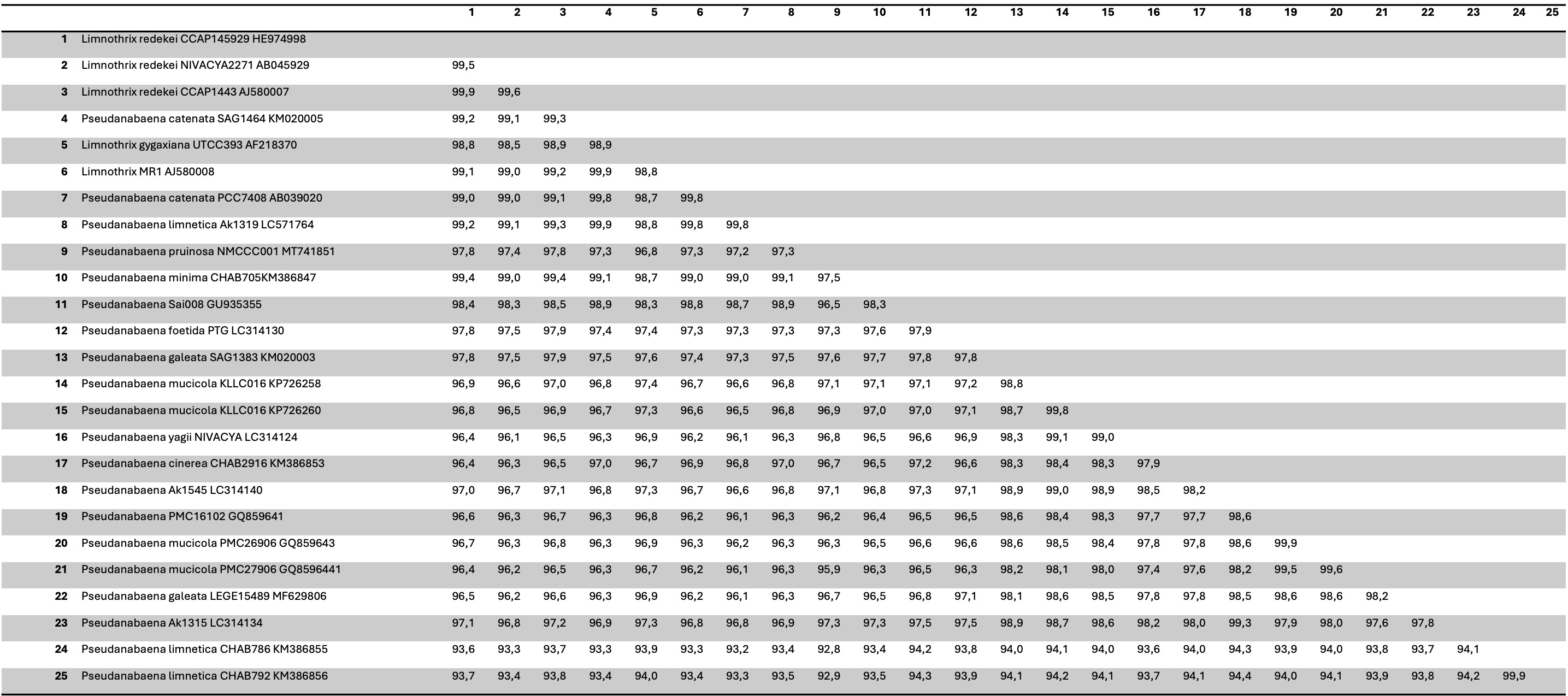
